# IFNγ-dependent metabolic reprogramming restrains an immature, pro-metastatic lymphatic state in melanoma

**DOI:** 10.1101/2024.12.02.626426

**Authors:** Triantafyllia Karakousi, Vanessa Cristaldi, Maria Luiza Lopes de Oliveira, Luiz Henrique Medeiros Geraldo, Tania J. González-Robles, Gabrielle da Silva, Alec P. Breazeale, Joel Encarnacion-Rosado, Joanna Pozniak, Alec C. Kimmelman, Kelly V. Ruggles, J. Chris Marine, Navdeep S. Chandel, Amanda W. Lund

## Abstract

Lymphatic vessels play a crucial role in activating anti-tumor immune surveillance but also contribute to metastasis and systemic tumor progression. Whether distinct lymphatic phenotypes exist that govern the switch between immunity and metastasis remains unclear. Here we reveal that cytotoxic immunity normalizes lymphatic function and uncouples immune and metastatic potential. We find that in mice and humans, intratumoral lymphatic vessel density negatively correlates with productive cytotoxic immune responses and identify IFNγ as an intrinsic inhibitor of lymphangiogenesis. Specific deletion of the *Ifngr1* in lymphatic endothelial cells (LECs) greatly expanded the intratumoral lymphatic network and drove the emergence of a tip-like endothelial state, promoting lymph node metastasis but not dendritic cell migration. IFNγ inhibits oxidative phosphorylation, which is required for proliferation and acquisition of the pathologic transcriptional state. Our data indicate that IFNγ induces a phenotypic switch in tumor-associated lymphatic vessels to reinforce canonical immune surveillance and block metastasis.

## Introduction

Lymphatic vessels regulate fluid homeostasis, lipid transport, immune surveillance, and waste clearance^1^. While adult dermal lymphatic vessels are largely quiescent, they often expand in a process termed tumor-associated lymphangiogenesis^2^. Increased lymphatic vessel density in the primary tumor is linked to increased lymph node (LN) metastasis, which is associated with poor prognosis in multiple solid tumors including melanoma^3,4^. Recent studies, however, demonstrated that vascular endothelial growth factor C (VEGF-C)-mediated lymphangiogenesis also leads to improved responses to immunotherapy in preclinical models of melanoma and glioblatoma^5,6^. This is presumed to be due to increased antigen transport and dendritic cell (DC) trafficking, which mobilizes antigen-specific adaptive immune responses and enhances anti-tumor T cell priming^6^. The therapeutic potential of a VEGF-C targeted approach is nevertheless limited by the induction of regional LN metastasis^7–9^, which can limit ongoing and future adaptive immune responses^10,11^. Therefore, targeting lymphangiogenesis in the clinical setting hinges on a deeper mechanistic understanding of how lymphatic vessels facilitate metastasis and immunogenicity for decoupling these processes.

While lymphatic vessels are indeed critical for the efficient initiation of adaptive tumor immune responses in both malignant and non-malignant settings^12,13^, new lymphatic growth, lymphangiogenesis, seems to be dispensable for the formation of a robust adaptive immune response against non-tumor insults in the skin^12^. Instead, activation of the pre-existing lymphatic vessel capillaries, which included the tightening of interendothelial junctions, was sufficient to boost DC migration from the infected skin to the draining LN and promote the early expansion of anti-viral CD8^+^ T cells^14^. Indeed, lymphatic endothelial cells (LEC) are activated by inflammatory mediators and fluid flows to increase expression of chemokines and adhesion molecules^12,15,16^ that may facilitate active transport and thereby tune more productive immune responses. These observations together support a hypothesis that it is the activation state of the peritumoral lymphatic vasculature, rather than absolute number, that determines their functional impact within the tumor microenvironment (TME).

In this study, we aimed to understand whether tumor-associated lymphangiogenesis was sufficient and necessary to boost adaptive immune surveillance in melanoma. Our data revealed a surprising negative correlation between intratumoral lymphatic vessel abundance and the accumulation of cytotoxic immunity in mouse and human melanomas. We were therefore motivated to understand the causal extrinsic drivers that regulate tumor-associated lymphangiogenesis as a function of local cytotoxicity and identified IFNγ as a potent, endogenous anti-lymphangiogenic cue that acted intrinsically to restrain LEC proliferation. The expanded intratumoral lymphatic vasculature that emerged in the absence of IFNγ-restrain was notably immature, displaying tip-like features that benefited the metastatic tumor cell but limited the migration of professional antigen presenting cells (APC). Importantly we find that IFNγ acts through metabolic reprogramming to reinforce the mature, quiescent LEC state and limit intratumoral expansion. Our data therefore implicate IFNγ as a critical differentiation cue that functions to normalize tumor-associated lymphatic vessel function to promote tumor immune surveillance while enforcing a barrier to metastasis.

## Results

### Lymphatic vessel density is inversely linked to antitumor immune responses

Lymphatic vessel density is associated with LN metastasis, yet boosting lymphatic vessel numbers through overexpression of VEGFC leads to better immunosurveillance and immunotherapy responses in preclinical melanoma and brain tumor models^5,6^. Still, lymphangiogenesis is not needed to establish potent immune responses during infection with the Vaccinia virus^14^. We therefore asked whether high lymphatic vessels density was necessary for potent anti-tumor immunity by evaluating the extent of intratumoral lymphatic involvement as a function of changes in intrinsic tumor immunogenicity.

We first used the isogenic YUMM1.7 and YUMMER1.7 murine melanoma cell lines (*Braf*^V600E^;*Pten*^fl/fl^;*Cdkn2a*^-/-^) that differ by their number of somatic mutations and thereby their response to immunotherapy^17,18^. As was previously reported^18,19^, YUMM1.7 tumors give rise to cold tumors with a myeloid bias, while YUMMER1.7 cells form hot microenvironments characterized by intratumoral cytotoxic CD8^+^ T cells producing IFNγ (**Figures S1A-C**). Rather than seeing a positive association between lymphatic density and the local CD8^+^ T cell response, we observed a significant reduction in intratumoral lymphatic vessel density (**Figures 1A** and **B**), lymphatic endothelial cell (LEC; CD45^-^CD31^+^gp38^+^) abundance (**Figures 1C** and **D**), and proliferation (**Figure 1E**) in the YUMMER1.7, hot tumor microenvironment. Conversely, when we quantified lymphangiogenesis in the autochthonous and poorly immunogenic *Tyr:Cre*^ERT^^2^;*Braf*^V600E^;*Pten*^fl/fl^ melanoma model^20^, we observed a progressive increase in lymphatic vessel density (**Figure 1F** and **G**) associated with a very low abundance of CD8^+^ T cells and a strong myeloid bias (**Figure S1F**).

**Figure 1.**
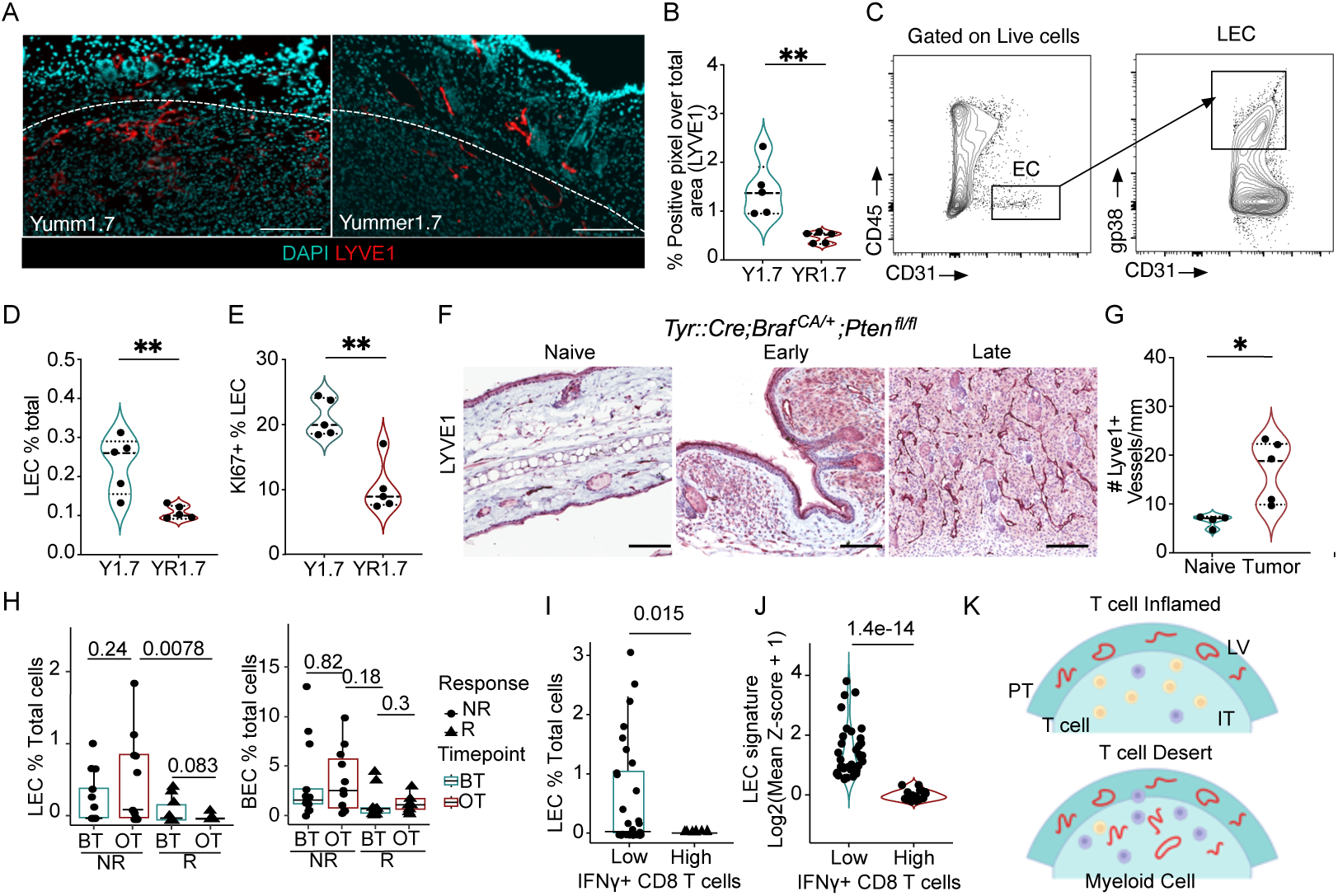
Cytotoxic tumor microenvironments are associated with reduced lymphatic vessel density. A. Representative immunofluorescence images of LYVE-1^+^ (red) lymphatic vessels from Y1.7 and YR1.7 tumors, nuclei are stained with DAPI (cyan), at day 12 of tumor growth. Scale bar = 100μm. B. Quantification of lymphatic vessel density from (A). C. Gating strategy to identify lymphatic endothelial cells (LEC; CD45^-^CD31^+^gp38^+^). D. LECs as a percentage of total live cells and E. Ki67 as a percentage of total LECs in Y1.7 and YR1.7 tumors. F. Representative IHC images of Lyve1 staining in naïve and Tyr::Cre;*Braf* ^CA/+^;*Pten* ^fl/lf^ tumors, G. Quantification of lymphatic vessels in naïve and Tyr::Cre;*Braf* ^CA/+^;*Pten* ^fl/lf^ tumors, each dot is a mouse n=4-5, H. Quantification of LECs and BECs as a percentage of total cells per melanoma patient in responders (R) and non-responders (NR) before and on immunotherapy from the single-cell dataset. I. Quantification of LECs in IFNγ^+^CD8 T low and high patient samples from the single-cell dataset, each point is a patient. J. Quantification of LEC signature in IFNγ^+^CD8^+^ T low and high patient samples from the TCGA dataset, each point is a patient. K. Schematic of peritumoral (PT) and intratumoral (IT) lymphatic vessels and T cells. Two-sided, unpaired Student’s t-tests (B-J), Wilcoxon rank test (H). *p<0.05; **p<0.001.

To determine if this inverse correlation between lymphatic vessels and an intratumoral cytotoxic CD8^+^ T cell response was also seen in human melanomas, we analyzed two separate datasets of melanoma patients, a single-cell human dataset from melanoma patients both before and during immunotherapy treatment^21^, and the cutaneous melanoma TCGA dataset^22^. In the single-cell dataset, LEC were annotated based on the expression of the lymphatic endothelial genes (*LYVE1, FLT4, PROX1, CCL21*) (**Figure S1G and S1H**), and were found to be more abundant in non-responder patients (**Figure 1H**), and inversely associated with the presence of *IFNG*-expressing *CD8A*^+^ T cells (**Figure 1I**). None of the other stromal components including the blood endothelial cells (BEC) (**Figure 1H**), cancer-associated fibroblasts (CAF) or pericytes exhibited these same shifts in abundance (**Figure S1I**), indicating a unique effect on LECs. Similarly, a negative correlation between *IFNG*-expressing *CD8A*^+^ T cells and LECs was observed in the TCGA dataset, where we compared the expression of a LEC signature score (*LYVE1, FLT4, PROX1, CCL21*) and the expression of cytotoxic T cell genes (*IFNG, CD8A*) (**Figure 1J and S1J**). Collectively, these data indicate that productive anti-tumor immune responses are inversely correlated with significant tumor-associated lymphatic expansion (**Figure 1K**), which indicates that lymphangiogenesis, in the setting of a poorly immunogenic tumor, is not sufficient to sensitize for response to immunotherapy.

### IFNγ suppresses tumor-associated lymphangiogenesis

These findings suggested the possibility that cytotoxic microenvironments might negatively regulate lymphatic expansion, which would be consistent with prior work demonstrating a role for IFNγ in patterning LN lymphatic networks^23^ and its lymphangiostatic function *in vitro*^24^. Consistently, analysis of canonical lymphangiogenic growth factors, revealed only a moderate increase in *Vegfd* expression (**Figures S2A**) with no observed change in *Vegfa* and *Vegfc* between the YUMM1.7 and YUMMER1.7 tumors. We therefore sought to determine if the elevated abundance of IFNγ^+^ CD8^+^ T cells (**Figure 1B-D**) and *Ifnγ* transcript (**Figure S2B**) seen in the YUMMER1.7 tumors could provide an endogenous, negative signal to intrinsically restrain tumor-associated lymphangiogenesis. We generated *Prox1:Cre^ERT^*^2^*;Ifnγr1^fl/fl^*mice, to specifically and inducibly knock out the *Ifnγr1* in adult LECs *in vivo*^25^. Adult mice were induced and rested one week prior to intradermal tumor implantation and the loss of IFNγ responsiveness specific to LECs was confirmed *ex vivo*^25^. To ask if loss of IFNγ-sensitivity was sufficient to rescue intratumoral lymphangiogenesis in immunogenic tumors, we intradermally transplanted YUMMER1.7 tumors into *Ifnγr1^iProx^*^1^ mice and *Ifnγr1*^WT^ littermate controls and collected the tumors 12 days post implantation. We found a striking rescue of intratumoral lymphangiogenesis evidenced by an increase in LYVE1^+^ pixel area (**Figures 2A** and **B**) and a trend in increase of LEC (**Figure 2C**) but not BEC numbers in the tumor microenvironment (**Figure 2D**). Strikingly, loss of the *Ifnγr1* on LECs resulted in intratumoral lymphatic expansion even in the presence of supraphysiological expression of the growth factor VEGFC by B16F10 melanoma cells also expressing the model antigen ovalbumin (OVA; **Figures 2E-H**). LECs isolated from B16-OVA-VEGFC and YUMMER1.7 tumors from *Ifnγr1^iProx^*^1^ mice were more proliferative based on Ki67 compared to littermate Cre-controls, while BECs from the same tumors did not show a difference in frequency or proliferation (**Figures 2I-K**). Similar results were seen in C57Bl/6 wildtype hosts with a neutralizing antibody against IFNγ, which increased the proliferation of both LECs and BECs in B16-OVA-VEGFC tumors (**Figures S2C**). Consistent with this *in vivo* data, IFNγ treatment limited the proliferation of human dermal LEC (HDLEC) *in vitro* (**Figure S2D**), forcing them to stall in the S phase of the cell cycle (**Figures S2E-F**). These data indicated that IFNγ is a potent inhibitor of proliferative lymphangiogenesis in the tumor microenvironment.

**Figure 2.**
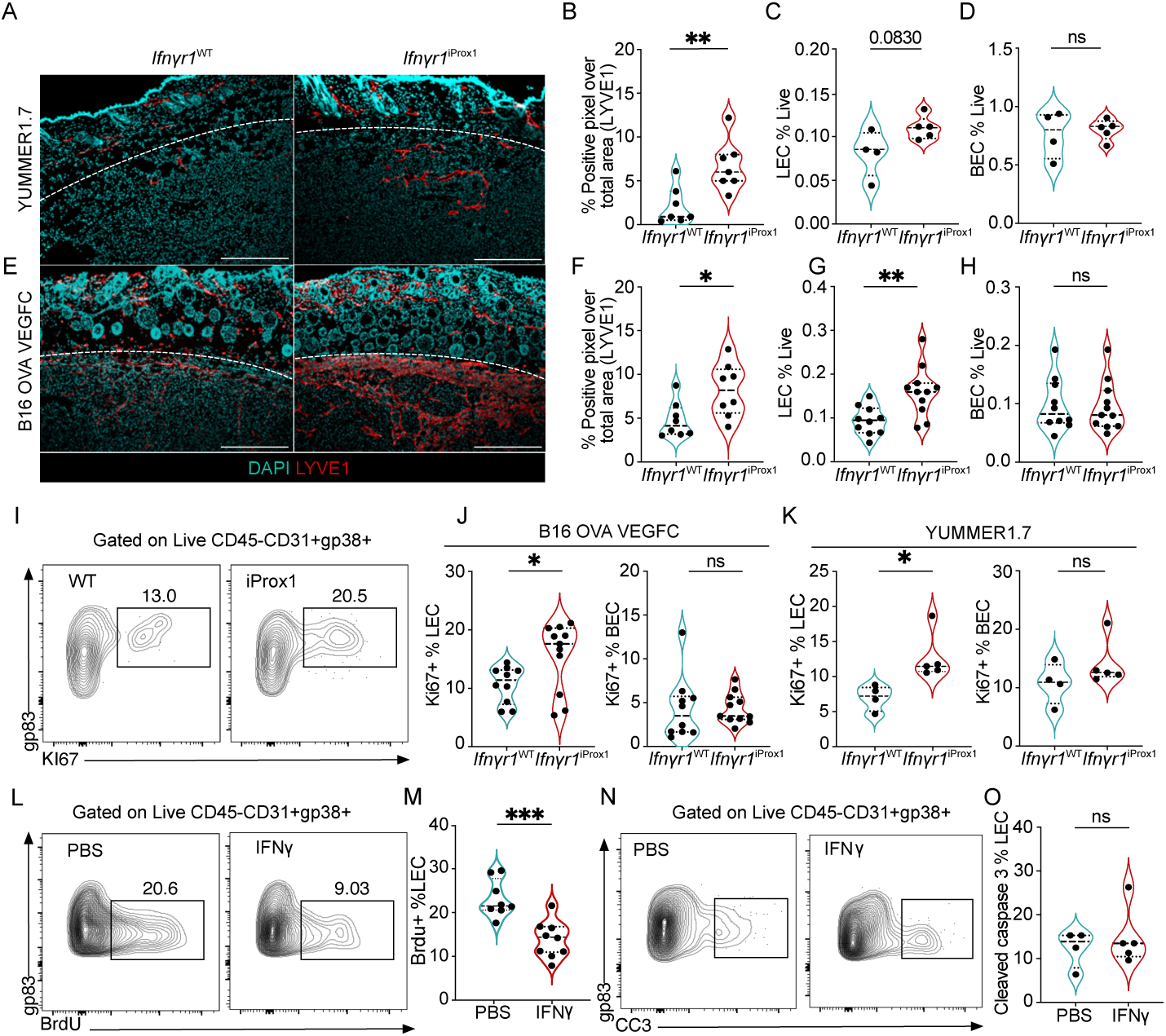
IFNγ restrains tumor-associated lymphangiogenesis. A. Representative immunofluorescence images of LYVE-1^+^ (red) lymphatic vessels and nuclei (DAPI, cyan), from YUMMER1.7 tumors grown in *Ifnγr1*^WT^ and *Ifnγr1*^iProx^^1^ mice, at day 12 of tumor growth. Scale bar = 500μm. B. Quantification of lymphatic vessel density (LVD) from image (A), each point represents a mouse (n=7). C. Lymphatic (LEC; CD45^+^CD31^+^gp38^+^) and D. blood endothelial cells (BEC; CD45^-^CD31^+^gp38^-^) as the frequency of total live cells from YUMMER1.7 tumors in the *Ifnγr1*^WT^ and *Ifnγr1*^iProx^^1^ mice, each point represents a mouse (n=4-5). E. Representative immunofluorescence images of LYVE-1^+^ (red) lymphatic vessels and nuclei with DAPI (cyan), from B16-OVA-VEGFC tumors grown in the *Ifnγr1*^WT^ and *Ifnγr1*^iProx^^1^ mice, at day 15 of tumor growth. Scale bar = 500μm. F. Quantification of LVD from (E), each point represents a mouse, (n=8). G. Quantification of LECs and H. BECs as the frequency of live cells from B16-OVA-VEGFC tumors in the *Ifnγr1*^WT^ and *Ifnγr1*^iProx^^1^ mice, each point represents a mouse, (n=10-11). I. Representative flow plots and J. quantification of Ki67 as a percent of LECs and BECs in B16-OVA-VEGFC tumors, (n=10-11) and K. YUMMER1.7 tumors from *Ifnγr1*^WT^ and *Ifnγr1*^iProx^^1^ mice, (n=2-5), each point is a mouse. L. Representative flow plots and M. quantification of percent of BrdU^+^ LECs in B16-OVA-VEGFC tumors from *Rag1*^-/-^ mice, treated with intratumoral injections of PBS or recombinant IFNγ, each point represents a mouse, (n=8-9). N. Representative flow plots and O. quantification of percent of cleaved caspase 3 (CC3) positive LECs in B16-OVA-VEGFC tumors from *Rag1*^-/-^ mice, treated with intratumoral injections of PBS or recombinant IFNγ as the frequency of total LECs, each point represents a mouse, (n=4-5). *p<0.05; **p<0.01; ***p<0.001. Unpaired, student’s t test.

In response to IFNγ, LECs upregulate immune inhibitory and antigen presentation molecules like PDL1 and MHCII which can contribute to the dampening of effector T cell responses^25,26,27^. A recent study using the B16-OVA-VEGFC tumor model argued that IFNγ induces OVA cross-presentation by LECs increasing their sensitivity to killing by cytotoxic TCR transgenic OTI T cells^28^. Our data, however, seemed to indicate an intrinsic change in proliferative potential. To determine if IFNγ puts a cell-intrinsic block on the expansion of LECs in the tumor microenvironment independent of cytotoxic T cells, we implanted B16-OVA-VEGFC tumors in Rag1^-/-^ mice, which lack T and B cells. At day 10 of tumor growth, we started intratumoral injections of mouse recombinant IFNγ or PBS for five consecutive days paired with the administration of Bromodeoxyuridine (BrdU) one day before euthanizing the animals. Tumor-associated LECs treated with IFNγ demonstrated reduced BrdU incorporation as compared to control, indicative of decreased proliferation upon IFNγ administration (**Figure 2L** and **M**). As previously reported^28^, there was no difference in apoptosis as evidenced by cleaved caspase 3 (CC3) staining (**Figures 2 N and O**). These data, therefore, argue that IFNγ inhibits LEC proliferation and is thereby a potent cell-intrinsic negative regulator of tumor-associated lymphangiogenesis.

### IFNγ enforces a differentiated LEC state in tumors

To begin to understand how IFNγ shaped the landscape of the tumor-associated lymphatic vasculature we performed single-cell RNA sequencing on live, CD45^-^CD31^+^ endothelial cells sorted from three *Ifnγr1*^iProx^^1^ and three *Ifnγr1*^WT^ B16-OVA-VEGFC tumors. Transcriptional analysis and visualization using uniform manifold approximation projection (UMAP) (**Figure S3A**), identified three LEC (*Prox1, Lyve1, Ccl21a, Pdpn*) and two BEC (*Pecam1, Aqp1*) clusters (**Figure S3B**). From the merged object, LECs subclustered into 4 unique capillary subsets all expressing *Lyve1*, *Prox1*, *Ccl21a, and Pdpn* and lacking expression of collector genes (e.g. *Foxc2*) (**Figure 3A** and **B**). Clusters 3 and 1 were transcriptionally less active, with high expression of genes involved in cholesterol homeostasis (*Fabp4, Fabp5*) typical of a more quiescent LEC state^29–31^. Cluster 2 expressed markers of proliferation including *Mki67* and *Top2a,* while cluster 0 activated genes related to cellular movement and Notch signaling. Interestingly, rather than seeing an increase in all 4 clusters in the absence of IFNγ signaling, we rather observed the selective expansion of cluster 0 in *Ifnγr1 ^iProx^*^1^ tumors. Cluster 0 was almost absent in the *Ifnγr1^WT^* LECs but made up almost 50% of the LECs from the *Ifn*γ*r1*^iProx^ group (**Figure 3C**). Interestingly, while *Ifnγr2* was uniformly expressed in all clusters, *Ifnγr1,* which encodes for the ligand binding component of the receptor ^32,33^, was specifically expressed in the wildtype cluster 0, perhaps indicating differential sensitivity to IFNγ amongst LECs in tumors (**Figure S3C**).

**Figure 3.**
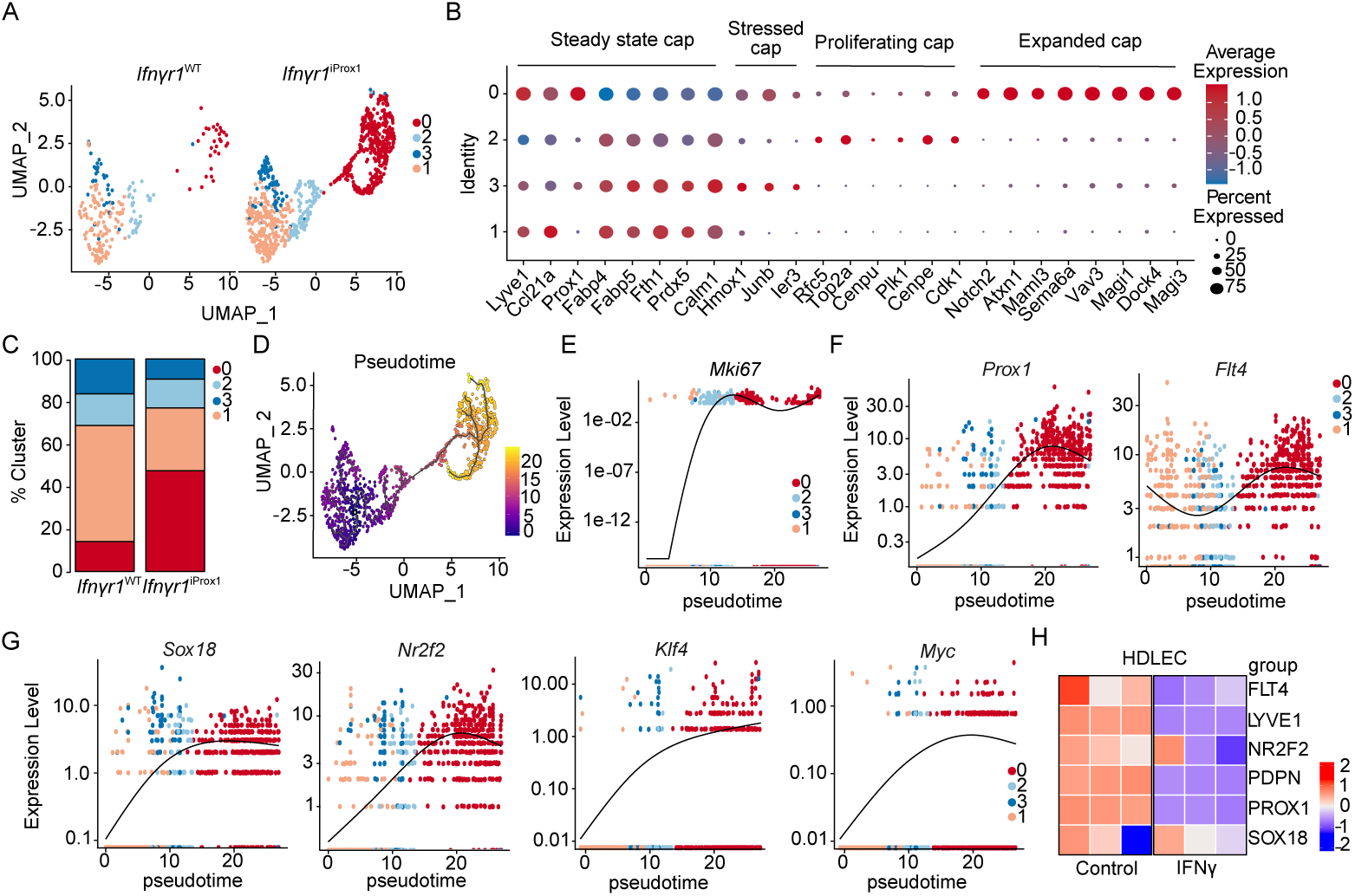
IFNγ promotes lymphatic differentiation away from an immature, tip-like state. A. Split UMAP scRNAseq analysis of tumor-associated LECs (LEC; *Prox1*^+^) sorted from B16-OVA-VEGFC tumors in *Ifnγr1*^WT^ and *Ifnγr1*^iProx^^1^ mice from. B. Average expression of marker genes for each cluster, normalized from -1 to 1. Size indicates percentage of cells in the cluster expressing the transcript. C. Cluster proportion in the *Ifnγr1*^WT^ and *Ifnγr1*^iProx^^1^ mice. D. Monocle 3 pseudotime analysis of the combined LEC object from (A). E-G. Gene expression as a function of pseudotime from (D). H. Heatmap of LEC fate genes expressed by human dermal LECs (HDLEC) treated with PBS or IFNγ (n=3, technical replicates) by bulk RNA sequencing.

These data seemed to indicate that a loss of IFNγ-restraint on proliferation allowed for the acquisition of a new LEC state that might have differential function in the tumor microenvironment. To understand the lineage relationship of the newly expanded LEC cluster with the rest of the LEC clusters, we performed pseudotime analysis using Monocle 3, selecting the quiescent cluster 1 as the beginning of the trajectory. One major trajectory arose from the analysis, consistent with a transition from clusters 1 and 3 through the proliferative cluster 2 into the expanded cluster 0 (**Figure 3D**). Adult dermal LECs are largely quiescent, terminally differentiated, and do not depend on VEGF-C–VEGFR-3 signaling for their maintenance^34^, but can be activated in tumors to re-establish a growth program^35^. Interestingly, analysis of transcripts that defined this growth trajectory in our data revealed upregulation of *Mki67* in the transition cluster, cluster 2 (**Figure 3E**), followed by a relative increase in expression of the LEC-fate defining transcriptional factor *Prox1* and growth factor receptor, *Flt4,* while the rest of the lymphatic markers like *Pdpn*, *Lyve1*, *Ccl21a* were constitutively expressed across clusters (**Figures 3F and S3D**). Aligned with the increase in expression of developmentally relevant lymphatic transcription factors and growth receptors, we also observed a progressive upregulation of transcription factors that regulate developmental lymphangiogenesis including *Sox18*^36^*, Nr2f2*^37^*, Klf4*^38^*, and Myc*^39^, indicating the possibility that LEC expansion in tumors requires dedifferentiation into a developmental LEC state (**Figure 3G**) and that IFNγ acts to maintain a mature and quiescent LEC population. IFNγ was sufficient to directly downregulated the expression of *Prox1*, *Flt4,* and the expression of *Nr2f2* and *Sox18* in HDLEC *in vitro* (**Figure 3H**), indicating that IFNγ directly suppresses the transcriptional activation of this immature LEC state.

### IFNγ inhibits an immature, tip-like state that promotes metastasis but not canonical tumor immune surveillance

The analyses presented above indicated that loss of IFNγ signaling in tumor-associated lymphatic vessels released the expansion of a pathological lymphatic state that appeared to activate transcriptional programs required for lymphatic development. We hypothesized that this pathological state might have distinct function from the pre-existing lymphatic vasculature on the probability of metastasis and immune surveillance. We found that B16-OVA-VEGFC tumors implanted intradermal in *Ifnγr1^iProx^*^1^ and *Ifnγr1^WT^*controls demonstrated equal rates of primary tumor growth (**Figure 4A** and **B**), but a significant increase in the dissemination of tumor cells to the tumor-draining LN as evidenced by the presence of LN pigmentation (**Figures 4C-D**). Interestingly, however, despite the dramatic increase in intratumoral lymphatic vessel density (**Figure 2**) and this increase in metastatic potential, we saw no significant difference in the transport of Evan’s blue injected intratumoral to the draining LN (**Figures 4E**), indicating that these pathologically expanded vessels were pro-metastatic but not canonically functional.

**Figure 4.**
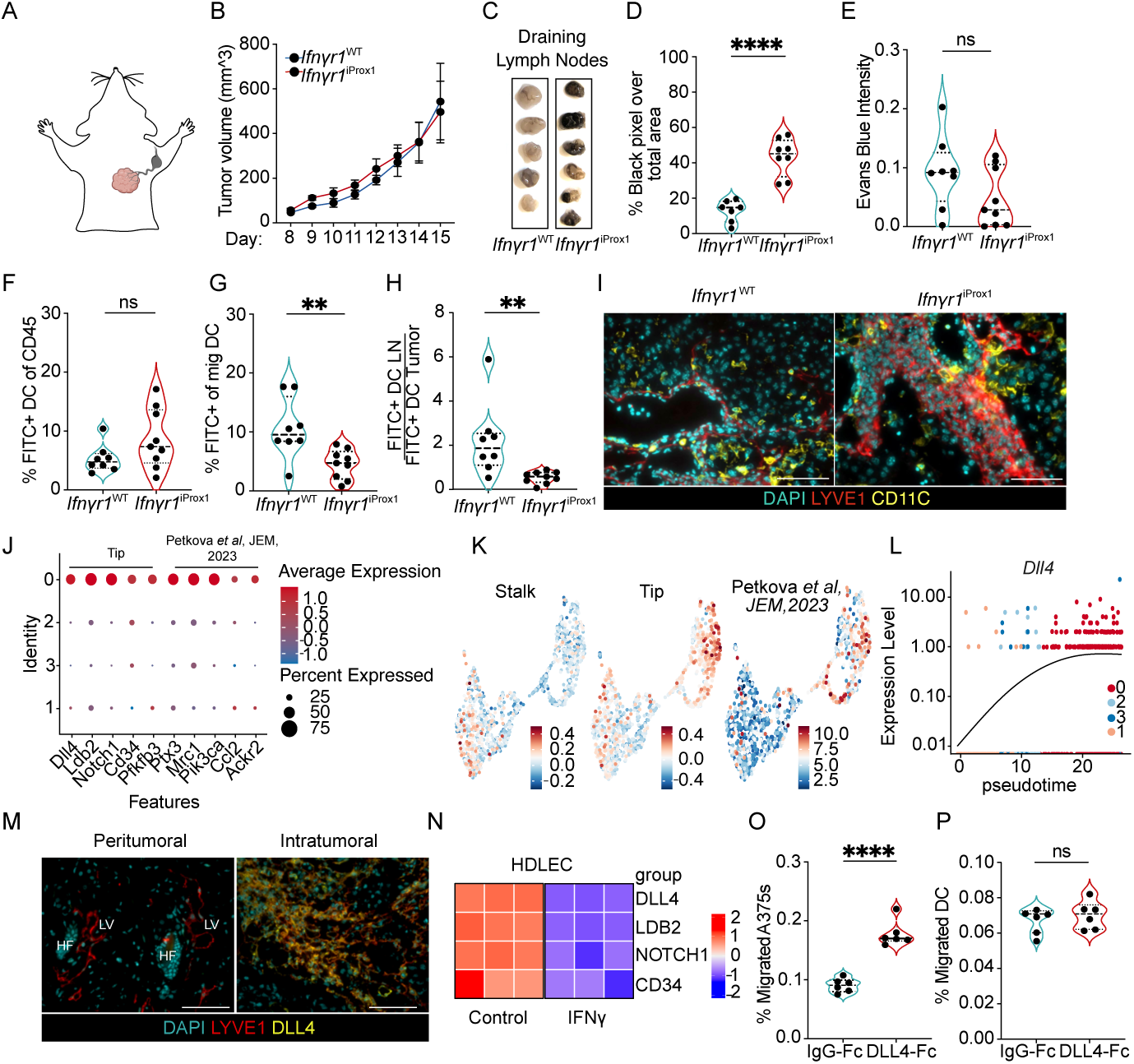
IFNγ-signaling in tumor-associated lymphatic vessels restrains a tip-like state that promotes lymph node metastasis but inhibits dendritic cell migration. A. Schematic of lymphatic drainage of the intradermal tumors on the top right side of the back. B. B16-OVA- VEGFC tumor growth (intradermal, 5x10^5^ cells) in *Ifnγr1*^WT^ and *Ifnγr1*^iProx^^1^ mice (n=5). C. Representative gross images of B16-OVA-VEGFC tumor-draining LNs from *Ifnγr1*^WT^ and *Ifnγr1*^iProx^^1^ mice. D. Quantification of black intensity over total LN area, each point represents a mouse, (n=7-8). E. Quantification of Evans blue intensity in B16-OVA-VEGFC tumor-draining LNs from *Ifnγr1*^WT^ and *Ifnγr1*^iProx^^1^ mice, each point represents a mouse, (n=8-9). F. Quantification of CD11C^+^MHCII^high^FITC^+^ migratory dendritic cells (mDC) as percent of live CD45^+^ cells in B16-OVA-VEGFC tumors and G. in tumor-draining LNs from *Ifnγr1*^WT^ and *Ifnγr1*^iProx^^1^ mice, each point represents a mouse, (n=8-9). H. Rate of migration as a ratio of FITC^+^CD11C^+^MHCII^high^ mDC in the LN to the tumor. I. Representative images of B16-OVA-VEGFC tumors from *Ifnγr1*^WT^ and *Ifnγr1*^iProx^^1^ mice. Lymphatic vessels (LYVE-1; red), DCs (CD11c; yellow, and nuclei (DAPI; cyan). Scale bars=100mm. J. Dot plot of select genes expressed in cluster 0, normalized from -1 to 1. Size indicates the percentage of cells in the cluster expressing the transcript. K. Stalk, tip, and Petkova et al module scores projected on combined UMAP. L. *Dll4* expression as a function of pseudotime. M. Representative immunofluorescence images of B16-OVA-VEGFC tumors from *Ifnγr1*^iProx^^1^ mice. Lymphatic vessels (LYVE-1, red); DLL4 (yellow); nuclei (DAPI, cyan). HF=Hair follicle, LV=Lymphatic vessel, scale bar = 100μm. N. Heatmap of tip-like genes expressed by HDLECs treated with PBS or IFNγ (n=3 technical replicates). O. Quantification of the percentage A375 melanoma, each point represents a technical replicate across two independent experiments, or P. bone marrow-derived dendritic cells (DC) that migrate across a transwell in the presence (DLL4-Fc) or absence (IgG-Fc) of adsorbed DLL4, each point represents a BDMCs from a single mouse, across two independent experiments. ****p<0.0001. Student’s t-test. *p<0.05; **p<0.01; ****p<0.0001.

While the increase in tumor dissemination to the LN we observed was consistent with prior work that demonstrated that intratumoral lymphatic vessel density and VEGFC positively associates with rates of LN metastasis in both mouse and human, the literature would also predict a concomitant increase in DC migration and therefore tumor immune surveillance. DC migration to the draining LN is critical for the generation of tumor-specific T-cell responses and tumor control^40^ and lymphatic vessels express the canonical chemoattractant, CCL21, which is required for optimal DC migration to LNs^40–42^. The presumed benefit to VEGF-C-induced lymphatic expansion in the context of immunotherapy is at least in part the increased rates of DC migration for *de novo* priming in the tumor-draining LN^5,6,43^. Surprisingly, however, despite the extensive expansion of the lymphatic vasculature, and the ability of the tumor cells to exploit this network for metastasis, the rate of DC migration on day 12 of B16-OVA-VEGFC tumor growth (**Figure S3E**), was not only not increased in *Ifnγr1*^iProx^^1^ mice, but rather decreased (**Figures 4F-H**). The reduction in DC migration we observed did not seem to depend on a failure of DCs to home to these expanded lymphatic vessels, which retained high expression of *Ccl21a* (**Figure S3E**). Indeed, we found abundant CD11c^+^ cells adjacent to intratumoral lymphatic vessels in both *Ifnγr1*^WT^ and *Ifnγr1*^iProx^^1^ tumors (**Figure 4I**), which may indicate that the chaotic, intratumoral endothelial structures that emerge in the absence of IFNγ signaling do not set up the gradients and adhesive structures optimally utilized by mature DCs for transendothelial migration and tissue egress. Importantly, these data indicate that this pathological, expanded lymphatic state may uncouple tumor and DC migration, providing an opportunity to explore the mechanisms that govern the functional impact of this state for future therapeutic intervention.

To therefore returned to the single cell to identify functional programs within the expanded, immature lymphatic cluster that might explain this differential function. We found that cluster 0 was further defined by high expression of endothelial tip cell marker genes like *Dll4, Cd34, Pfkfb3, and Ldb2*, as well as *Ptx3, Mrc1*, and *Pi3kca* (**Figure 4J**), which define a newly characterized immune interacting cluster of LECs at the tips of lymphatic vessels in a mouse model of oncogene-driven lymphatic malformation^17^. A signature previously established for tip cells^45^ as well as for the immune interacting Ptx3^+^ LEC signature^44^ enriched in the expanded LECs while a stalk cell signature enriched in the rest of the clusters (**Figure 4K**). DLL4/NOTCH1 exhibits both pro- and anti-lymphangiogenic features in lymphatic vessels, which likely reflects its role in specification of the tip-like state^46,47^. While inhibition of DLL4 induces increased ectopic sprouting during lymphatic development^48^, postnatal lymphangiogenesis, in the context of wound healing^49^ and regeneration of the intestinal lacteal^47^, depends on DLL4 expression where it promotes lymphangiogenesis by enhancing VEGF-C/VEGFR3 signaling^50^. Interestingly, in the adult, DLL4 expression is inhibited in lumenized lymphatic vessels by laminar shear stresses^50^, which simultaneously imposes quiescence, a hallmark of dermal lymphatic vessels. Similar to the fate-determining, developmental transcription factors, the pseudotime trajectory analysis revealed that *Dll4* expression was progressively enriched in the expanded LEC cluster (**Figure 4L**). We therefore sought to use DLL4 as a marker of the expanded cell state and through immunofluorescence analysis confirm the expression of DLL4 on the expanded, intratumoral but not pre-existing dermal LECs in B16-OVA-VEGFC tumors from *Ifnγr1*^iProx^^1^ mice (**Figure 4M**). Importantly, again, IFNγ downregulated the expression of the tip cell markers in HDLECs *in vitro*, and thereby directly inhibited entry into the tip-like state (**Figure 4N**). Interestingly, multiple markers of the expanded cell state are implicated in lymphatic metastasis, including *Ptx3*, *Ccl21a*, and *Ccl2*^51–53^, suggesting that the altered surface and chemokine repertoire of these intratumoral lymphatic vessels may preferentially support tumor cell migration, DLL4 is also directly implicated in metastasis in both blood^54–56^ and lymphatic vessels^57^. DLL4 expressed by LECs initiates a positive feedback loop with melanoma cells via NOTCH3 to increase their metastatic potential^57^. Consistently, we found that DLL4-Fc, but not IgG-Fc, when coated on transwell plates, increased the migration of A375 human melanoma cells but had no effect on mouse DCs migrating towards a CCL21 gradient (**Figures 4O** and **P**). These data indicated that IFNγ not only negatively regulates the proliferation of tumor-associated lymphatic vessels, but actively restrains a pathological phenotypic state that, likely through multiple mechanisms, benefits invasive tumor cells but stalls the migration of activated DCs.

### IFNγ selectively blocks the expansion of the immature tip-like lymphatic endothelial capillary subset by rewiring mitochondrial metabolism

Our data above point to an IFNγ-dependent mechanism that not only restrains the proliferation of LECs within the TME but specifically restricts an immature, tip-like, pathological lymphatic state while leaving pre-existing lymphatic vessels largely intact. We therefore sought to define the molecular mechanisms that drove this effect to nominate new therapeutic targets for lymphatic modulation in tumors. To identify IFNγ-dependent pathways that might limit the expansion of the immature tip-like LEC state we performed pathway analysis of differentially expressed genes in the expanded, tip-like cluster (cluster 0) between *Ifnγr1^iProx^*^1^ and *Ifnγr1^WT^*tumors. We found an upregulation in pathways related to mitochondrial metabolism including oxidative phosphorylation (OXPHOS), reactive oxygen species (ROS), and hypoxia in the absence of IFNγ-signaling (**Figure 5A**). Scoring clusters with a signature of mitochondria OXPHOS complexes revealed low expression in the tip-like cluster 0, which was partially rescued in the *Ifnγr1^iProx^*^1^ compared to the *Ifnγr1^WT^* (**Figure 5B**). Mitochondria respiration is pivotal for the development and maintenance of the lymphatic vasculature during embryonic development, with LEC-specific deletion of mitochondrial complex III leading to a failure of lymphatic vessel development in multiple organs including the skin, the gut, and heart ^58^. We, therefore, hypothesized that IFNγ might limit LEC proliferation by targeting mitochondria respiration. Consistent with an increase in OXPHOS, we observed increased mitochondrial ROS, which is generated as a by-product of active mitochondrial respiration, in tumor-associated LECs from *Ifnγr1*^iProx^^1^ mice bearing day 12 B16-OVA-VEGFC tumors (**Figures 5C** and **D**). To directly test the hypothesis that IFNγ is sufficient to inhibit OXPHOS, we treated HDLEC *in vitro* with IFNγ or PBS for 3 days. Consistent with the *in vivo* results, IFNγ-directly decreased mitochondria ROS production in HDLEC (**Figures 5E** and **F**) and the ability of HDLEC to produce energy through OXPHOS as both basal and maximal oxygen consumption rates (OCR) were decreased (**Figures 5G** and **H**). Furthermore, consistent with the transcriptional changes (figure 5B), western blot analysis of HDLEC lysates treated with IFNγ demonstrated the downregulation of mitochondrial complexes III-UQCRC2 and II-SDHB further supporting a direct blockade of OXPHOS by IFNγ in HDLECs through direct effects on the expression of complexes of the mitochondrial respiratory chain (**Figures S4**). Intriguingly, in addition to its role in regulating LEC proliferation, OXPHOS is required to maintain the developmental program that drives lymphatic patterning during embryogenesis by transcriptionally controlling the expression of key lymphatic genes like *Prox1* and *Flt4*^58^. Indeed, direct inhibition of mitochondrial complex III with antimycin-a, phenocopies the effect of IFNγ on HDLEC proliferation and transcriptional state, decreasing the expression key lymphatic transcripts^58^ as well as the tip-like signature (**Figure 5I**).

**Figure 5.**
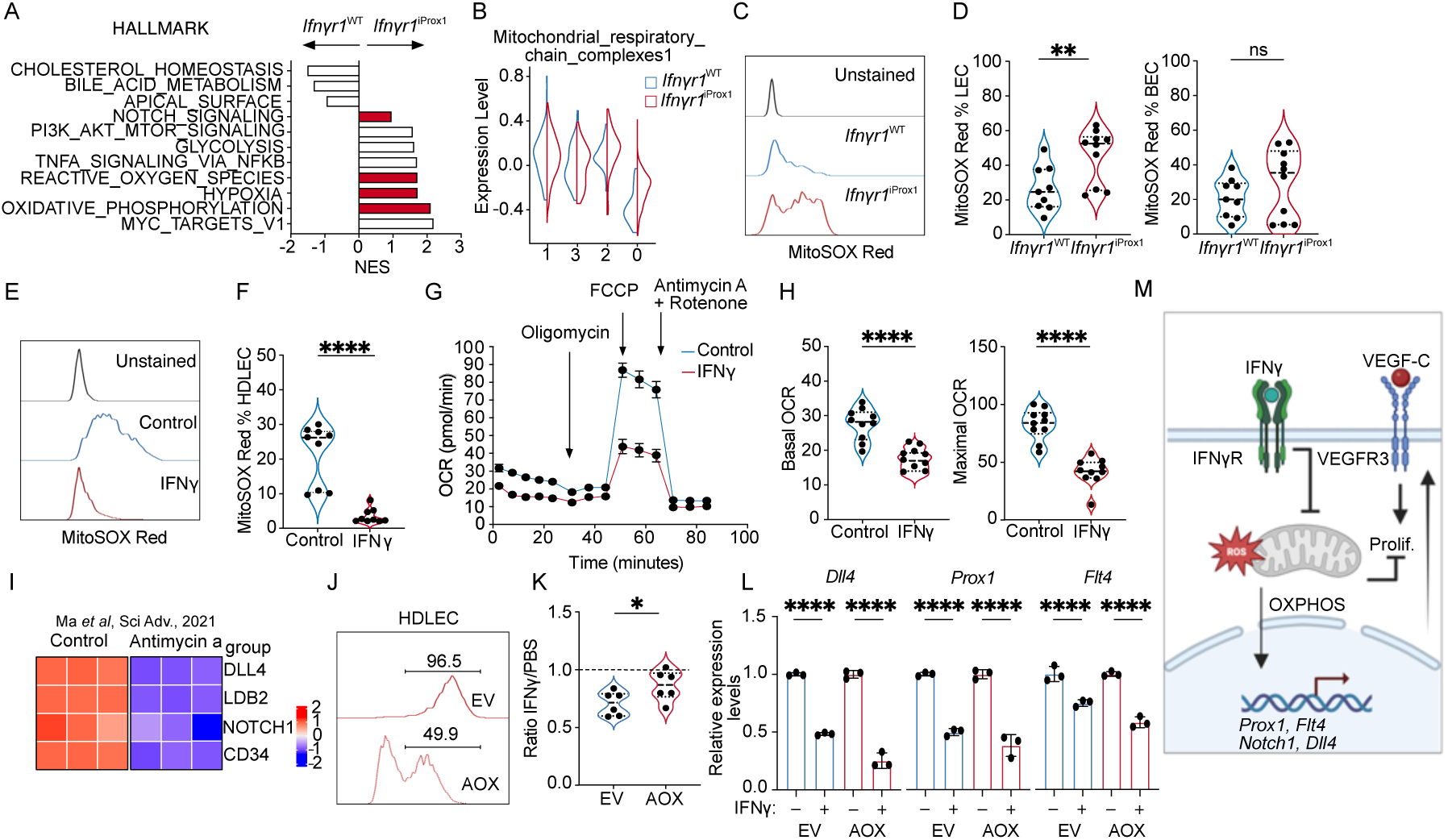
IFNγ inhibits proliferation through effects on mitochondrial respiration. A. Pathway analysis of differentially expressed genes in cluster 0 between *Ifnγr1*^WT^ and *Ifnγr1*^iProx^^1^ LECs from Fig. 4. B. Mitochondrial respiratory chain complex module score as a function of cluster and experimental group. C. Representative histogram of Mitosox Red staining in tumor-associated LECs (LEC; CD45^-^CD31^+^gp38^+^) from B16-OVA-VEGFC tumors from *Ifnγr1*^WT^ and *Ifnγr1*^iProx^^1^ mice. D. Mitosox Red quantification in LEC and blood endothelial cells (BEC; CD45^-^ CD31^+^gp38^-^) from B16-OVA-VEGFC tumors from *Ifnγr1*^WT^ and *Ifnγr1*^iProx^^1^ mice, each point represents a mouse, (n=9-10). E. Representative histogram and F. quantification of Mitosox Red in HDLEC treated with PBS or IFNγ (72 hrs). G. Seahorse analysis of IFNγ-conditioned (48hrs) HDLECs relative to PBS-treated controls. H. Basal and maximal oxygen consumption rate (OCR) from (E). I. Heatmap of tip-like genes from bulk RNA sequencing data of HDLECs treated with PBS or antimycin-a scaled and reanalyzed (Ma et al Sci Adv 2021). J. Representative histogram of HDLECs transduced with empty vehicle control (EV-RFP) or a plasmid expressing the alternative oxidase (AOX-RFP). K. Quantification of RFP^+^ EV and AOX HDLEC numbers after treatment with IFNγ (72 hrs), presented relative to PBS controls. L. *Dll4*, *Prox1, Flt4* expression in sorted EV-RFP or AOX-RFP transduced HDLECs after IFNγ or PBS treatment. M. Schematic illustration of the role of IFNγ on proliferation and transcriptional control of selected genes. *p<0.05, **p<0.01, ***p<0.001, ****p<0.001.

To determine whether the impact of IFNγ on proliferation and transcriptional state depended on its effects on OXPHOS, we rescued complex III activity in the presence of IFNγ. To do so, we transduced HDLECs with lenti-viruses expressing the alternative oxidase enzyme (AOX) or empty vector controls both carrying the red fluorescent protein (RFP). AOX bypasses the activity of complex III and IV and restores electron transport chain (ETC) activity, and at the same time uncouples ETC from mitochondria ROS production^59,60^. After HDLEC transduction with either the EV-RFP control or the AOX-RFP lenti-viruses (**Figure 5J**) we treated them with IFNγ or PBS control for 3 days and assessed their relative growth. AOX expression rescued the IFNγ mediated proliferation defect in HDLECs (**Figure 5K**), but interestingly, did not restore the *Dll4*, *Prox1* and *Flt4* transcription (**Figure 5L**), indicating that while electron flux is required for proliferation, mitochondrial ROS, which is generated by the activity of complex III and can leak through the mitochondria intermembrane space to the cytosol^61,62^ to act as signaling intermediates^63–65^, may be required to induce the developmental transcriptional state. These mechanistic data therefore indicate that IFNγ simultaneously places a block on proliferation and developmental transcriptional programs by directly inhibiting OXPHOS (**Figure 5M**). These data collectively indicate that cytotoxic immunity, and specifically IFNγ, reinforce a mature, differentiated LEC state through metabolic reprogramming that acts as a barrier to regional tumor progression and promotes immune surveillance.

### The immature LEC state stratifies melanoma patients for worse survival and poor immunotherapy response

Consistent with our findings, using single cell analyses a recent study^66^ independently characterized the presence of a tip-like LEC state in human solid tumors, which was found to be mutually exclusive with an antigen presenting (apLEC) state (**Figure 6A**). We therefore asked whether the human tip-like LECs identified in this study shared the same immature gene expression program with the expanded cluster in our single cell data. Analysis of gene expression as a function of pseudotime trajectory revealed a progressive enrichment of the embryonic developmental LEC factors (**Figure 6B**) in the tip-like LEC trajectory, which were mutually exclusive with the expression of IFNγ-related antigen presentation molecules as well as the *IFNGR1* transcript (**Figure 6C**). These data confirm that the transcriptional states we observe in mice are not only present but are consistent with a model where IFNγ is a key switch driving specification along these two distinct tumor-associated fate trajectories.

**Figure 6.**
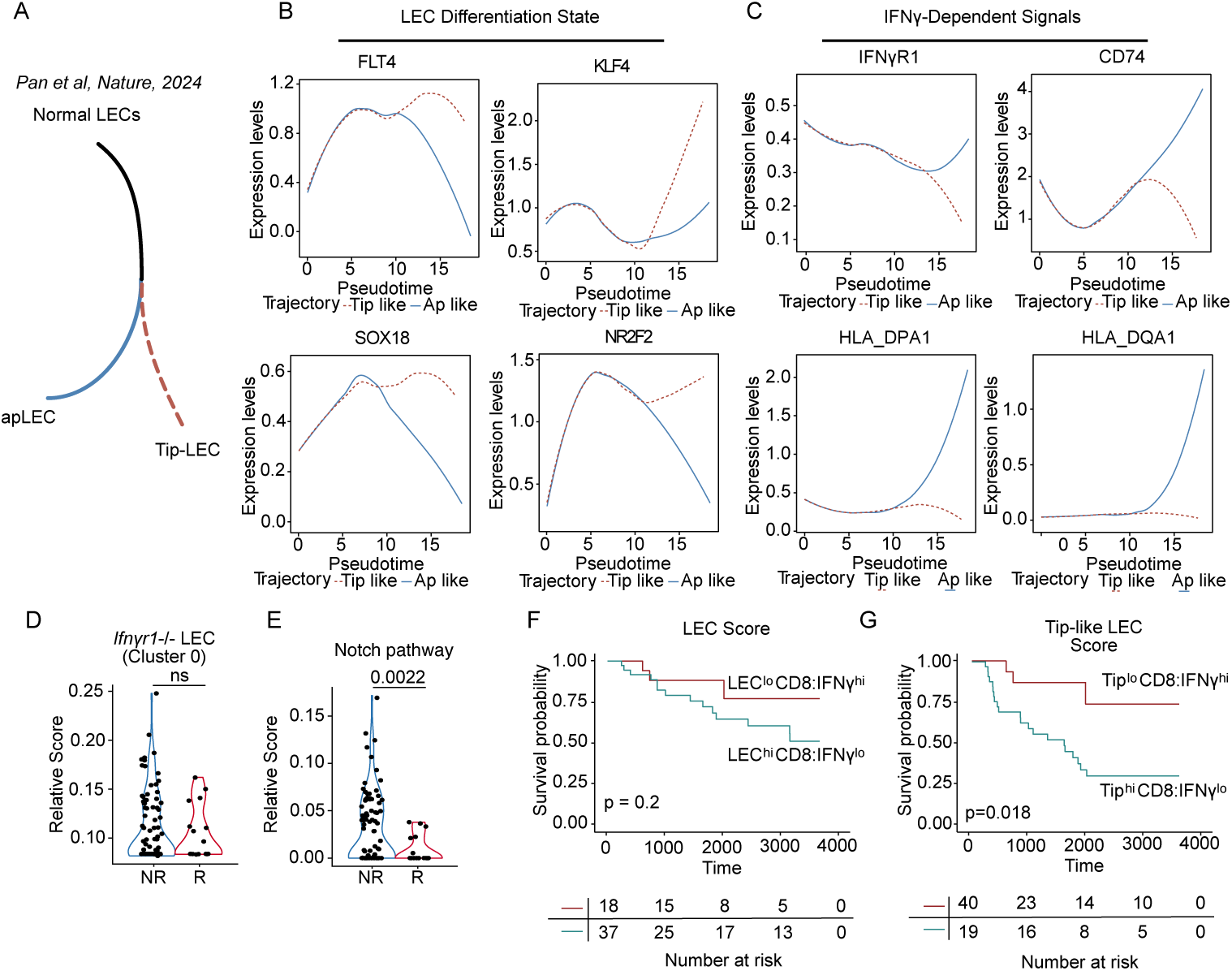
The tip-like lymphatic state is associated with poor response to immunotherapy and worse overall survival in human melanoma. Α. Schematic illustration of the antigen presenting (apLEC) and the tip (Tip-LEC) trajectories from Pan *et al*, Nature, 2024. B-C. Pseudotime trajectories of select genes in the Pan *et al*, Nature, 2024, dataset. D. Module score of LEC cluster 0 from *Ifnγr1*^iProx^^1^, and E. Notch pathways signatures on non-responder (NR) and responder (R) melanoma patients before and on immunotherapy from the single-cell dataset. Each point is a single cell. F. Kaplan-Meier curve depicting overall survival of cutaneous melanoma patients from the TCGA-SKCM dataset, stratified with LEC^hi^CD8:IFNγ^lo^ and LEC^lo^CD8:IFNγ^hi^ and G. the Tip-like LEC cluster signatures Tip^hi^CD8:IFNγ^lo^ and Tip^lo^CD8:IFNγ^hi^, (p-values shown in log-rank test).

To further this idea, we returned to the scRNAseq data collected from metastatic melanoma patients treated with ICB^21^, where we had initially observed the inverse correlation between LEC and CD8^+^ T cell abundance (**Figures 1H** and **I**). Here, we generated a signature from our expanded cluster 0 (**Figure 6D**) as well as genes curated from the Notch pathway (**Figure 6E**), and found indeed that LECs in patients responding to therapy, with a higher abundance of *IFNG*^+^*CD8*^+^ T cells^21^, significantly suppressed our pathological LEC state relative to LECs found in non-responders. These data provide striking evidence for the conservation of transcriptional states between mouse and human and their regulation by cytotoxic immunity, and particularly IFNγ.

Finally, we wanted to understand the extent to which the specific lymphatic state, rather than simply the number of lymphatic vessels, could contribute to outcome in patients. We turned to the TCGA data set where we had found that a generic LEC signature was mutually exclusive with CD8A and IFNγ expression (**Figure 1L and S1K**). When we stratified patients according to these two groups, LEC^hi^CD8:IFNγ^lo^ and LEC^lo^CD8:IFNγ^hi^ (**Figure S1K**) and looked at overall survival in a cutaneous melanoma cohort, we found no statistical difference in survival probability (**Figure 6F**), indicating that LEC abundance itself was not sufficient to select patients for progression. However, when we modified the LEC signature to include markers of the tip-like state we now saw a significant stratification of survival probability (**Figure 6G**). These data indicate that patients that have mobilized significant CD8^+^ T cell responses, sufficient to reprogram the transcriptional state of the intratumoral LEC, may see better outcomes, importantly demonstrating that the intratumoral LEC state is likely of more significance than their absolute abundance.

## Discussion

During the early stages of tumor progression, the lymphatic vasculatures provide the necessary route for tumor immune surveillance and thereby can contribute to tumor control^13^. Overtime, however, cooption of the lymphatic vasculature by a developing tumor leads to LN metastasis, which itself can install mechanisms of immune suppression that further increase the probability of systemic disease^10,11^. We recently outlined the hypothesis that a phenotypic switch in lymphatic function must underlie this conversion from immune surveillance to metastasis^2^. Here we present data to indicate that IFNγ promotes a mature lymphatic state that reinforces the barrier to metastasis and promotes immune surveillance. Mechanistically, we demonstrate that IFNγ directly limits OXPHOS, which is required for the acquisition of a developmental and tip-cell program that directly fuels lymphatic growth and provides a functional scaffold for tumor cell invasion. These data support a model whereby an immunogenic tumor capable of mobilizing early cytotoxic immune responses might be less likely to metastasize, and indeed microsatellite instable gastric cancers exhibit reduced LN metastasis relative to microsatellite stable disease^67,68^. Our findings reveal lymphatic heterogeneity in tumors and underscore the delicate balance required to maintain a functional network of activated lymphatic vessels—crucial for sustaining anti-tumor immunity—while preventing the rise of dysfunctional LEC states that favor tumor progression.

Lymphatic vessels are necessary for the induction of adaptive immune responses following peripheral tissue challenge and tumor development^12,13^. Consistent with a causal association between lymphatic transport and anti-tumor immune surveillance, several studies have reported a positive correlation between VEGFC^6^ or lymphatic vessel density^13,25,69^ and tumor-infiltrating lymphocytes. Paradoxically, however, while VEGFC/VEGFR3 enhances DC trafficking and *de novo* T cell priming in the draining LN^5,7^, increased VEGFC also generates immunosuppressive microenvironments^7^ and is drives LN metastasis in mice^9^. Therefore, while VEGFC/VEGFR3 signaling seems to enhance immunotherapy responses in both melanoma^6^ and brain cancer^5^ mouse models, this perceived immune benefit is likely to be confounded overtime by LN metastasis^10,11^, and the pleiotropic roles of VEGFC in the local tissue microenvironment complicate a mechanistic understanding for the specific impact of lymphatic growth on the therapeutic impact. Here we demonstrate that the mobilization of productive, cytotoxic CD8 T cell responses provide negative feedback on lymphatic growth and serves to reinforce their maturation even in the presence of ectopic VEGFC expression. Our analyses in human datesets^21,70^ also demonstrate a clear mutual exclusivity between cytotoxic CD8 T cells and LECs and consistent with this we recently demonstrated that intratumoral T cell infiltrates, likely indicating more cytotoxic activity, were associated with reduced lymphatic vessel density in human cutaneous primaries^71^. These data therefore support the interpretation that increased lymphatic vessel density is not required for the generation of potent anti-tumor immunity and, in fact, the converse, that highly cytotoxic immune microenvironments are associated with reduced local lymphatic growth.

The negative association between cytotoxic T-cell infiltrates and lymphatic density indicated the possibility of direct negative feedback. We found that IFNγ produced by cytotoxic CD8 T cells inhibits tumor lymphangiogenesis and tumor progression by intrinsically inhibiting the proliferation and, therefore, expansion of new lymphatic vessels. These data are consistent with the observations that LN lymphangiogenesis is enhanced and more persistent in mice lacking T cells and IFNγ^23^ and *in vitro* studies clearly demonstrate a direct impact of IFNγ on lymphatic proliferation and sprouting^24^. In contrast, a recent study proposed that IFNγ activates exogenous antigen presentation by LECs and thereby renders them sensitive to killing by activated TCR transgenic T cells^28^. Indeed, peripheral tissue lymphatic vessels are potent sensors of IFNγ and we previously demonstrated that the level of surface PD-L1 and MHCII was finely tuned by the abundance of antigen-reactive CD8 T cells and IFNγ. Highly immunogenic tumors, e.g. YUMMER1.7, therefore activated an IFNγ-dependent immune regulatory program that limited effector T cell responses and tumor control^25^. We saw similar immunoregulatory function in the setting of viral infection where IFNγ levels increase dramatically between days 7-15 post infection in concert with productive, anti-viral T cell infiltrates^25^. It may therefore be that the impact of IFNγ on the lymphatic vasculature depends on a fine-tuning of the magnitude and persistence of its expression within tissue. Indeed, excessive pruning of the blood vasculature, which can be mediated by IFNγ^72^, leads to hypoxia and immune suppression^73^. Of note, we do not observe significant expression of PDL1 or antigen presentation molecules in intratumoral LECs in our model, which may indicate that the physiological levels of IFNγ in a progressing tumor prune lymphangiogenesis through intrinsic, anti-proliferative effects without activating immune suppression pathways.

Not only did IFNγ limit LEC proliferation, but it also appeared to serve as a potent differentiation cue in the TME. IFNγ sensing inhibited the emergence of an immature, tip-like LEC state associated with high expression of factors that regulate the growth and development of the lymphatic vasculature including *Prox1*^74^, *Nr2f2*^37^, *Sox18*^36^, *Myc*^39^, *Klf4*^38^*, and Dll4*^49,50^. This transcriptional program thereby appeared to render the intratumoral LECs more sensitive to VEGFC/VEGFR3 signaling driving expansive and invasive lymphatic growth. Indeed, PROX1 and VEGFR3 work in a positive feedback loop to maintain a LEC state that is necessary for LEC specification and early development^58^. The immature lymphatic cluster, was also transcriptionally similar to a recently identified LEC subtype which arose at the tips of lymphatic vessels in the skin of a *Pik3ca-*driven lymphatic malformation^44^, again linking this developmental state to aggressive and dysregulated peripheral tissue lymphatic growth. Of note, this embryonic transcription factor signature was also found in human LECs in a pan-cancer single-cell analysis^66^, where it was mutually exclusive with an antigen-presenting LEC subtype that bore signs of an IFNγ signature^66^. Our data therefore indicate that IFNγ acts as an endogenous differentiation cue for the lymphatic vasculature restraining their expansion in the presence of potentially tumor-promoting inflammation, and thereby limiting metastatic progression. Interestingly, IFNγ seems to act broadly to reinforce differentiation in diverse cell types including stem-like T cells^75^, dendritic cells^76^, and monocytes^77,78^, and may therefore act as a global determinant of peripheral tissue immune homeostasis.

We posited that an understanding of the IFNγ-dependent mechanisms that regulate both lymphatic growth and maturation could lead to new therapeutic strategies for lymphatic modulation in tumors. Our data indicate that IFNγ directly inhibits mitochondrial metabolism to both limit proliferation and promote their differentiation in tumors. While no studies have elucidated the metabolic requirements for lymphatic growth in tumors, developmental lymphangiogenesis depends upon fatty acid oxidation (FAO)^31^ and glycolysis^39^. FAO is necessary for the generation of acetyl-coA which is used by histone acetyltransferase p300 to acetylate lymphatic genes, providing a mechanism through which metabolic changes could impact LEC state. Consistent with our data, mitochondrial respiration reinforces the PROX1/VEGFR3 feedback loop and is required for developmental lymphangiogenesis^58^. Direct inhibition of complex III *in vitro* is sufficient to reduce expression of *Prox1*, *Flt4*, and *Nrp2*^58^, a phenotype that we find to be consistent with IFNγ treatment. Indeed, our data indicate that IFNγ directly reduces complex III, which leads to impaired proliferation and restriction of the developmental state. We propose that while this *Prox1*^high^ state is necessary for the specification of LEC from the venous endothelium during the earliest stages of lymphatic development, restriction of this program in adult skin, particularly in the context of inflammation, may be necessary to maintain mature vessels and thereby promote their canonical function.

These data indicate that IFNγ may act to tune LEC state through its effects on mitochondrial respiration and associated mitochondrial metabolites^79,80^. Bypassing complex III with AOX to restore electron flux was sufficient to rescue proliferation in the context of IFNγ treatment but notably did not restore expression of some of the key developmental transcripts, e.g. *Prox1* and *Dll4*. AOX does not restore mitochondrial ROS production, which can control transcriptional programs through Hif1α stabilization^81,82^, and may indicate that there are distinct determinants of lymphatic growth and fate downstream of mitochondrial respiration. Interestingly, this link between LEC metabolism and LEC fate is supported by additional studies where FAO^83,84^, ketone body oxidation^85^, and autophagy^86^ impact the acetylation of key lymphangiogenic genes and thereby lymphatic growth. IFNγ treatment^87^ and inhibition of autophagy^86^ both increase the accumulation of lipid droplets and impact mitochondrial respiration, perhaps indicating convergence of these pathways in the regulation of LEC state. These data again emphasize the potential for targeted metabolic pathways to tune lymphatic vessel function *in vivo* and open interesting questions as to how diet may impact tumor-associated lymphangiogenesis, tumor progression, and tumor immune surveillance.

VEGFR3-targeted therapies aimed at limiting lymphatic growth in the TME have to date had limited success^88,89^, perhaps requiring a more targeted approach to tuning lymphatic vessel function. Our data indicate that IFNγ, secreted by cytotoxic CD8 T cells, constrains the expansion of immature, tip-like lymphatic vessels with pro-metastatic features by blocking OXPHOS. This population bore similarity to the LEC state enriched in melanoma patients non-responsive to immunotherapy, associated with poor outcomes in the skin cutaneous melanoma dataset from the TCGA, and associated specifically with poor outcome in hepatocellular carcinoma^66^. We therefore propose that inhibiting OXPHOS in tumor-associated lymphatic vessels may be an effective strategy to flip the balance between metastatic and immunogenic LEC function. Key to the efficacy of such a strategy, however, will be the extent to which IFNγ or OXPHOS inhibitors can contract or normalize an already expanded tumor-associated lymphatic vasculature. IFNγ induces blood vessel normalization in the TME^90,91^ and therapeutic administration leads to vascular regression tumor ischemia and necrosis^72^, which may suggest a parallel effect could be had on tumor-associated lymphangiogenesis. Unlike with the blood vasculature, however, expanded lymphatic networks can persist in the absence of growth factor maintenance^92^, which may indicate resistance to extrinsic contraction cues. Therefore, whether metabolic targeting of the pathologic state will be an efficacious therapeutic strategy in established tumors remains to be carefully tested. In sum, we propose that IFNγ acts as an endogenous switch to prune the lymphatic vasculature, regulating the balance between growth and maturation. We therefore propose the concept of lymphatic vessel normalization as a therapeutic strategy to reinforce canonical, immunogenic lymphatic function in melanoma and other solid tumors.

## Methods

### Mice

C57BL/6J and Rag1^-/-^ mice were purchased from The Jackson Laboratory. Prox1-Cre-ER^T^^2^ mice^93^ were provided by V.H. Engelhard (University of Virginia, Charlottesville, VA) in agreement with T. Makinen (Uppsala University, Uppsala, Sweden). IFNγR1^fl/fl^ mice^94^ (stock no. 025394) and Tyr::Cre;*Braf* ^CA/+^;*Pten* ^fl/lf^ mice^20^ (stock no. 013590) mice were purchased from Jackson. The Prox1-Cre-ER^T^^2^; IFNγR1^fl/fl^ mice were created by crossing the Prox1-Cre-ER^T^^2^ with IFNγR1^fl/fl^. All breeding was maintained at NYU in specific pathogen-free facilities. All mice were previously backcrossed over 10 generations to the C57BL/6 background. Mice were housed in specific pathogen-free rooms under 12-hour light/12-hour dark cycles with an ambient temperature of approximately 70 °C and 50% humidity. Water and standard chow were provided ad libitum. For all in vivo studies, sex-matched 8-10-week-old mice were used with at least three to five mice per group. All animal procedures were approved by and performed in accordance with the Institutional Animal Care and Use Committee at NYU.

### Cell lines

The YUMM1.7 and YUMMER1.7 cell lines^17,18^ were provided by M. W. Bosenberg (Yale University), and passaged in 1:1 DMEM (Cytiva, SH30243.FS), F-12 supplemented with 1% l-glutamate (Thermo Scientific,11765054), 1% non-essential amino acids (Sigma-Aldrich, M7145), 10% FBS, and 1% penicillin-streptomycin (Lonza, BW17602E). Primary Human Dermal Lymphatic Endothelial Cells (HDLECs) were purchased from PromoCell and passaged according to the manufacturer’s recommendations (C-12216). The B16-OVA-VEGFC^7^ cells were passaged in DMEM High Glucose (Cytiva, SH30243.FS) supplemented with 10% FBS and 1% penicillin-streptomycin (Lonza, BW17602E). Recombinant huma IFNγ (PeproTech,300-02) was reconstituted in sterile PBS. A375 (ATCC, CRL-3224) were passaged in DMEM (ATCC, 30-2002) supplemented with 10% FBS, and 1% penicillin-streptomycin (Lonza, BW17602E).

### In vitro cell culture

For HDLEC proliferation, HDLECs were seeded in 6-well plates at 30K or 60K per well. The following day they were treated with PBS control or IFNγ (0.5ug/ml) in fresh media, and counted every day for 4 days using a cell counter. Treatment was replenished every day. To produce the AOX-RFP and EV-RFP lentiviruses, we transfected 293T cells (ATCC) with pMD2.G (Addgene,12259) and psPAX2(Addgene,12260) packaging vectors along with the expression plasmids (gifted by Dr. Navdeep Chandel) using Lipofectamine 3000 Transfection Reagent. The virus was collected and filtered through 0.45uM PVDF filter before being added to HDLECs. Three days after transduction HDLECs were plated at 30K per well in 12 well plates and treated with PBS or IFNγ for 3 days. Relative proliferation of the transduced cells was quantified using flow cytometry. For the qPCR analysis, RED AOX HDLECs or RED EV HDLECs treated with PBS or IFNγ for 3 days were sorted, and RNA was extracted using the RNeasy mini kit (Qiagen, 74104) according to the manufacturer’s protocol.

### BMDC generation

Mice were sacrificed and bone marrow was extracted from the long bones of the legs and pooled per mouse. 10M bone marrow cells were plated per well in tissue-culture-treated 6-well plates in 4 ml of complete RPMI medium supplemented with glutamine, 2-mercaptoethanol (Sigma-Aldrich, M6250), 10% FBS and 20 ng/ml GM-CSF (R&D Systems, 415-ML-010). Half of the medium was removed on day 2 and replenished with warmed complete RPMI medium with 40 ng/ml GM-CSF and 10 ng/ml IL-4 (Biolegend, 574304). The culture medium was entirely changed on day 3 and replaced by fresh warmed medium with 20 ng/ml GM-CSF and 5 ng/ml IL-4. On day 6 cells were harvested for phenotyping and experiments.

### Transwell assay using the Boyden chamber

Tumor cell migration: The membranes of Boyden chambers (Corning,3421) were coated with 100 ng/ml Dll4-Fc (Acro Biosystems, DL4-H5254) or IgG-Fc (Sino Biological, 10702-HNAH-100) overnight at 4°C. After 3 washes with sterile PBS, 30K A375s were seeded on the upper chamber of the Boyden chamber in serum-free media and 10% FBS-supplemented media was added to the bottom chamber. Cells were left to migrate through the 5um porous membrane for 8 hours. Membranes were swapped with a Q-Tip to remove the non-migrated cells on the top side of the membrane, excised, fixed with PFA, and stained with DAPI.

DC migration: The membranes of Boyden chambers were coated with 100 ng/ml Dll4-Fc or IgG-Fc overnight at 4°C. After 3 washes with sterile PBS, 30K LPS stimulated BDMCs were added to the upper chamber, 100 ng/ml CCL21 (R&D Systems, AF457) was added to the lower chamber and BMDCs were left to migrate for 3 hours. RPMI medium supplemented with glutamine, 2-mercaptoethanol, 10% FBS with 20 ng/ml GM-CSF, and 5 ng/ml IL-4 was used for the experiment. The upper chambers were removed and total numbers of migrated BMDCs were quantified with flow cytometry.

### qPCR analysis

RNA was harvested from tumor tissues using the RNeasy mini kit (Qiagen, 74104) according to the manufacturer’s protocol. cDNA was synthesized from RNA using the High-Capacity cDNA Reverse Transcription Kit (Thermo Fisher Scientific,4368814) according to the manufacturer’s protocol. Real-time PCR was performed with the StepOne Plus PCR system using the PowerUp SYBR Green Master Mix (Thermo Fisher Scientific, A25741). Primer sequences used were as follows:

Human-Prox1-F: 5’-CTGAAGACCTACTTCTCCGACG-3’;

Human-Prox1-R: 5’-GATGGCTTGACGTGCGTACTTC-3’;

Human-Flt4-F: 5’-TGCGAATACCTGTCCTACGATGC-3’;

Human-Flt4-R: 5’-CTTGTGGATGCCGAAAGCGGAG-3’;

Human-Dll4-F: 5’-CTGCGAGAAGAAAGTGGACAGG-3’;

Human-Dll4-R: 5’-ACAGTCGCTGACGTGGAGTTCA-3’;

Human-Gapdh-F: 5’-GTCTCCTCTGACTTCAACAGCG-3’;

Human-Gapdh-R: 5’-ACCACCCTGTTGCTGTAGCCAA-3’

Mouse-Ifnγ-F: 5’-CAGCAACAGCAAGGCGAAAAAGG-3’

Mouse-Ifnγ-R: 5’-TTTCCGCTTCCTGAGGCTGGAT-3’

Mouse-Vegfa-F: 5’-CTGCTGTAACGATGAAGCCCTG-3’

Mouse-Vegfa-R: 5’-GCTGTAGGAAGCTCATCTCTCC-3’

Mouse-Vegfd-F: 5’-CTCCACCAGATTTGCGGCAACT-3’

Mouse-Vegfd-R: 5’-ACTGGCGACTTCTACGCATGTC-3’

Mouse-Vegfc-F: 5’-CCTGAATCCTGGGAAATGTGCC-3’

Mouse-Vegfc-R: 5’-CGATTCGCACACGGTCTTCTGT-3’

Mouse-Actin-F: 5’-CATTGCTGACAGGATGCAGAAGG-3’

Mouse-Actin-R: 5’-TGCTGGAAGGTGGACAGTGAGG-3’

### Cell cycle analysis

To quantify cell cycle changes, HDLECs were seeded in 6 well plates at 60K per well. The following day they were treated with PBS control or IFNγ (0.5ug/ml) for 3 days as described. Following the cells were detached and processed with the Click-iT™ Plus EdU Alexa Fluor™ 488 Flow Cytometry Assay Kit (Thermo Scientific, C10632) per the manufacturer recommendations.

### Tumor studies

Mice were anesthetized using vaporized isofluorane (at 1-3% isofluorane; oxygen flow rate at 0.5-1.0 L/min) and 500K B16-OVA-VEGFC, YUMM1.7, or YUMMER1.7 tumor cells were implanted intradermally into the back of mice. Tumor growth was measured daily using digital calipers for the long and short axes. Average diameters were used to calculate spherical volume. For treatments, murine recombinant IFNγ (PeproTech, 315-05) was reconstituted in sterile saline and administered intratumorally daily for 5 days with a Hamilton syringe. Neutralizing IFNγ (200ug/ mouse; BioXCell, XMG1.2, BE0055) or IgG (200ug/mouse; BioXCell, HRPN, BE0088) were administered intraperitoneally every two days starting at day 5 of tumor growth. Prox1-Cre-ER^T^^2^; IFNγR1^fl/fl^ and littermate controls, were treated with tamoxifen (Sigma-Aldrich, T5648-1G) as recommended by the Jackson Laboratory, 2mg of tamoxifen (75mg/kg diluted in corn oil) for five consecutive days followed by one week of rest. All tumor experiments were conducted in accordance with IACUC guidelines. For the bromodeoxyuridine (BRDU) pulse experiments, mice received an initial dose of 2mg BRDU (Fisher Scientific, H2726006) in saline intraperitoneally and maintained on BRDU 0.8mgml^-1^ water until the day of the sacrifice.

### Fluorescein Isothiocyanate Painting

Mice were anesthetized as above, and the tumors were painted with 20 μl of a 5% FITC (Sigma-Aldrich, F4274-50MG) solution prepared in acetone. 24 hours later, mice were euthanized and tumors and draining lymph nodes were collected.

### Evans blue drainage

Mice were anesthetized as above and 10 ul of 5% Evans Blue (Sigma-Aldrich, E2129) solutions in PBS were injected intratumorally using a Hamilton syringe. 10 minutes later mice were euthanized, and the draining lymph nodes were collected. Lymph nodes were placed in a 200 ul formamide solution (Sigma-Aldrich, F9037) and incubated for 24 hours at 55°C. 100 ul were transferred in a flat bottom 96 well plate and absorbance was measured at 620nm.

### Flow cytometry and antibodies

Tumors were harvested and digested with 220 U ml^−1^ collagenase IV (Invitrogen, 17104019) and 80 U ml^−1^ DNASE I (Sigma-Aldrich, 04536282001) for 30 min at 37°C. Digests were then passed through a metallic mesh and a 70-μm pore filter. Lymph nodes were smashed and passed through 70-μm pore filter. For surface stains, single-cell suspensions were stained with antibodies in FACS buffer (1%BSA in PBS) for 30 minutes at 4°C, in the dark. Staining of Ki67 (SolA15) was performed with True-Nuclear™ Transcription Factor Buffer Set (Biolegend, 424401) as per manufacturer protocol. Samples were analyzed on a BD LSR II, or BD FACSymphony. FlowJo 10.8.1. was used to analyze flow cytometry data. Antibodies were purchased from BioLegend, Cell Signaling Technologies, Invitrogen, BD Biosciences, and Tonbo and included: Ghost Dye Red 780 for dead cell detection, CD8 (53-6.7), CD45 (30-F11), CD4 (GK1.5), NK1.1 (PK136), F4/80 (T45 2342), I-A/I-E (M5/114.15.2), CD11c (N418), CD11b (M1/70), Ly-6C (HK1.4), Ly-6G (1A8), CD31 (MEC13.3), gp38 (8.1.1), Cleaved Caspase 3 (Asp175). For the IFNγ (XMG1.2) staining mice were injected intravenously through the tail vein with 500 ul of 0.5 mg/mL brefeldin A (BFA) in PBS 6 hours before sacrifice and the staining was performed with the Fixation/Permeabilization Solution kit (BD, 554714).

### Immunofluorescence and immunohistochemical staining

Tumors and LNs were fixed in 2% paraformaldehyde for 24 hours at 4°C, transferred to 15% sucrose overnight at 4°C, followed by 30% sucrose overnight at 4°C. Following they were indirectly frozen in optimal cutting temperature compound OCT. 5-10μm sections were blocked using 2.5% BSA solution plus Triton X100 in PBS for 1 hour at room temperature. Primary antibodies were added overnight and incubated at 4°C. Sections were washed with PBS 3 times and incubated with secondary antibodies in 1.25% BSA, 0.1Triton X100 in PBS for 2 hours at room temperature. Slides were sealed with SlowFade Gold antifade reagent with DAPI (Invitrogen, S36936). Images were acquired with Kyance and processed with ImageJ. Antibodies were purchased from Reliatech, Thermo Scientific, Biolegend, Abcam and included: LYVE1 (103-PA50), CD8 (53-6.7), DLL4 (PA5-46974), CD11c (N418). Tyr::Cre;*Braf* ^CA/+^;*Pten* ^fl/lf^ mouse ears were resected and fixed with formalin for 24 hours, followed by dehydration with 70% ethanol and embedding in paraffin blocks. 5-10μm thick sections were deparaffinized and stained. Sections were bleached prior to staining in 10% H202 for 10 min at 65°C.

### Mitosox staining

For HDLEC: Cells treated with PBS or IFNγ for 3 days were collected and counted as per manufacturer guidelines (PromoCell Inc). 1M cells per condition were washed in 5 ml falcon tubes with HDLEC media (PromoCell Inc, C-22020) with no serum addition and stained with the Mitosox Red reagent (Thermo Scientific, M36008) for 20 min at 37°C. Following, cells were washed 3 times with HDLEC media with serum and the addition of DAPI for live dead detection and acquired at BD FACSymphony.

For tumors: Tumors were quickly collected and minced in room temperature RPMI (Cytiva, SH30027.FS) without serum and antibiotics. They were then passed through a metallic mesh and a 70 μm cell filter to acquire single-cell suspensions. The cells were then stained for 20 min with Mitosox Red and the flow antibodies at 37°C, washed 3 times with serum-added RPMI, and acquired at BD FACSymphony.

### Western blot

HDLECs were treated with PBS or IFNγ for 3 days in 6 well plates and washed with ice-cold PBS 1x. 100 microliters of ice-cold RIPA lysis buffer was added to each well and the cells were scrapped and transferred in 1.5 ml tubes and left on ice for 30 minutes followed by centrifugation at maximum speed at 4°C. Protein lysates were quantified with the BCA protein quantification kit. 30 ug of protein was loaded per well on a Surepage Bis-Tris 4-12% precast gel and run for 2.5 hours at 120 Volt. Proteins were then transferred on a methanol-activated PVDF membrane for 2 hours at 60 Volt. Following the membrane was blocked for 1 hour at RT with Every blot blocking buffer (Abcam, 12010020). The membrane was stained with the total OXPHOS antibody cocktail (Abcam, ab110411) overnight at 4°C, washed 3 times with Tris-buffered saline (TBST), and incubated with the secondary goat anti-mouse IgG Polyclonal Antibody (IRDye® 800CW) (LI-COR Biosciences, 926-68071) for 2 hours at RT. Similarly, the membrane was incubated overnight with a b-actin antibody (Cell signaling, 4967S), followed by TBST washes and incubation with the secondary goat anti-rabbit IgG polyclonal antibody (IRDye® 680RD) (LI-COR Biosciences, 926-68071). The protein signal was acquired with the Bio-rad ChemiDoc MP Imaging System. Band intensity was calculated with Image J.

### Seahorse

HDLECs were pre-treated with PBS or IFNγ for 2 days lifted using the DetachKit (PromoCell Inc, C41210) according to the manufacturer protocol, and re-plated in equal numbers in the 96 well plate Seahorse Xp (Agilent Technologies, 101085-004) at thirty thousand cells per well. The cells were left to attach overnight and maintained in PBS or IFNγ media. Once cells were fully attached their media was changed in Seahorse XF Assay Medium with a pH of 7.4 (Agilent Technologies, 103575-100),10 mM Glucose (Agilent Technologies, 103577-100), 1 mM Sodium Pyruvate (Thermo Scientific, 11360070) and 2 mM Glutamine (Agilent Technologies, 103579-100). HDLECs were maintained for 1 hour in a CO2-free incubator at 37 °C. Basal metabolism and mitochondrial stress test assay was performed as per manufacturer recommendations with final well concentrations of Oligomycin (Cayman Chemical Company, 11342) 2.5umol/L, FCCP (Cayman Chemical Company,15218) 2umol/L, and Antimycin/Rotenone (Sigma Aldrich, A8674/Cayman Chemical Company 13995) 0.5umol/L. Analysis was performed using Seahorse Wave Desktop Software (Agilent Technologies).

### Single-cell sequencing

B16-OVA-VEGFC were intradermally implanted in *Ifnγr1*^iProx^^1^ and *Ifnγr1*^WT^ mice. On day fifteen, tumors were collected and processed as above. Three tumors per condition were pooled and stained (DAPI^-^, CD31^+^, CD45^-^) and sorted separately. The sorted cellular suspensions (*Ifnγr1*^WT^ ;CRE- and *Ifnγr1*^iProx^^1^ ;CRE+) were loaded on a 10x Genomics Chromium instrument to generate single-cell gel beads in emulsions. Libraries were prepared using single cell 3′ reagent kits v3.1 (Chromium Next GEM Single Cell 3′ GEM, Library & Gel Bead Kit v3.1, 16 rxns PN-1000121; 10x Genomics) and were sequenced using Illumina Novaseq 6000. Both experiments used the transcriptome mm10-2020-A (mouse) and Cell Ranger 7.0.0. Before filtering, 2502 and 2491 cells were sequenced. Data were analyzed and visualized using Seurat packages in R. Quality control was performed on sequenced cells to filter out cells based on the number of RNA features (genes) (100 < genes < 10000), mitochondrial reads (<5%), ribosomal protein genes (20%) and ribosomal RNA genes (25%). LogNormalize was used to normalize for gene expression (scale factor, 10,000) and log-transform the results. ElbowPlot and JackStraw plots were used to determine the proper number of principal components.Cells were clustered using a shared nearest-neighbor modularity optimization–based clustering using FindNeighbors and FindClusters functions at a resolution of 0.3. Gene signatures were calculated using the AddModuleScore function in Seurat. We used Monocle 3^95^ for the analysis of our Seurat Object to infer cellular trajectories. A root node was established in Seurat cluster 0, which served as the starting point for the pseudotime analysis. Seurat’s object was then rearranged to allow for a more refined identification of cellular states.

scRNA-seq data from the human samples were acquired from Pozniak *et al* ^70^. The identification of CD8 T cells was performed as described previously^21^. Stromal cells were identified based on the genes signature acquired from Jerby-Arnon *et al*^96^ and then re-clustered obtaining LECs, BECs, Pericytes and CAFs. Identity of LECs was confirmed based on the expression of *LYVE1*, *FLT4*, *PROX1*, *CCL21*.

### Bulk RNA sequencing and gene set enrichment analysis

To analyze transcriptomic changes in the context of treatment with recombinant human IFNγ (n = 3) versus PBS Ctrl group (n = 3), samples were prepared for bulk mRNA sequencing following a polyA-enriched paired-end sequencing protocol (Illumina Truseq Stranded mRNA v2 protocol # 20020595) for the Illumina Novaseq X-Plus system platform. Analysis of bulk RNA sequencing was performed at NYULMC HPC UltraViolet (formerly BigPurple) cluster using software provided in the Seq-N-Slide pipeline (https://github.com/igordot/sns), through the *rna-star* followed by *rna-star-groups-dge* routes. Briefly, after quality control assessment with MultiQC (python/cpu/v2.7.15)^97^ and sequencing adaptor trimming with Trimmomatic (v0.36)^98^, reads were aligned to the mm10/GRCm38 mouse reference genome with a splice-aware (STAR v2.7.3a)^99^ alignment, followed by featureCounts (subread/v1.6.3)^100^ to generate the RNA counts table. These were then normalized for gene length and sequencing depth and tested for differential mRNA expression between IFNγ and Ctrl groups using negative binomial generalized linear models implemented by the DESeq2 v1.40.1^101^ R package (r/v4.1.2). RStudio v4.3.0 was used for additional downstream analyses and visualizations. Differential expression was assessed by principal component analysis (prcomp function from the stats v4.3.0 R package) or unsupervised hierarchical clustering (pheatmap v1.0.12 and ComplexHeatmap v2.16.0) and visualized by a volcano plot (EnhancedVolcano v1.18.0) and TPM expression heatmap illustrating genes of interest. Differentially expressed genes (DEGs) identified were further analyzed for gene set enrichment analysis (GSEA) and pathway analysis using R packages: fgsea v1.26.0 and msigdbr v7.5.1.

### Statistical analysis

Statistical analysis was performed using GraphPad Prism 10 software. Data were subjected to D’Agostino-Pearson, Kolmogorov-Smirnov, or Shapiro-Wilk tests to determine the distribution of data. Comparisons between two groups were conducted using unpaired Student’s t tests (parametric) when data were normally distributed or Mann-Whitney test and Wilcoxon matched-pairs signed-rank test (nonparametric). Comparisons between more than two groups that were normally distributed were conducted using one-way analysis of variance (ANOVA) with Tukey’s or Dunnet’s multiple comparisons test, whereas Kruskal-Wallis one-way ANOVA with Dunn’s multiple comparison test was used if one of the groups was not normally distributed. The proper sample size was based on prior experience. Statistical analysis of sequencing data was performed with pairwise Wilcoxon rank test and Bonferroni correction in R. P values are reported as follows: *P ≤ 0.05, **P ≤ 0.01, ***P ≤ 0.001, and ****P ≤ 0.0001. Error bars show the mean ± SEM.

## Resource Availability

Further information and requests for resources and reagents should be directed and will be fulfilled by the lead contact.

## Data and code availability

Data will be made publicly available upon publication.

## Acknowledgements

The authors acknowledge technical assistance and critical feedback from Alexandra Figueroa, Ekaterina Novikova, Molly Cohn, Devyon McDonnough and the rest of the Lund lab. This work was supported by grants from the Cancer Research Institute (Lloyd J. Old STAR Award to A.W.L.) and the NIH (R01CA238163, U54CA263001 to A.W.L.). T.K. is supported by an A.G. Leventis Foundation Fellowship. T.J.G.R. and K.V.R. are supported by funding from HHMI Gilliam Fellowship (GT15758) and T.J.G.R. received funding from the NIH (T32GM136542). J.P. received a Marie Curie Fellowship (H2020-MSCA-IF-2019, #896897) and Stichting tegen Kanker fellowship (2023/2310). J-C.M received funding from the VIB Grand Challenges Program (POINTILLISM) and FWO/FRS-FNRS (EOS, # 40007513), and received support from VIB core facilities (https://vib.be/en/technologies/core-facilities): VIB Single Cell Core, VIB Spatial Catalyst, FACS core.

## Author contributions

Conceptualization: TK, AWL

Methodology: TK, LHMG, TJGR, JER, JP, NC

Investigation: TK, VC, MLLdO, GdS,

APB Funding acquisition: AWL

Reagents: NC

Supervision: ACK, KVR, J-CM, AWL

Writing – original draft: TK, LHMG, AWL

Writing – review & editing: TK, VC, MLLdO, LHMG, TJGR, GdS, APB, JER, JP, ACK, KVR, J-CM, NSC, AWL

## Declaration of interests

A.W.L. reports consulting services for AGS Therapeutics. All other authors declare no conflicts of interest.

## Supplemental Information

Document S1. Figures S1-S4.

