## Supplemental Figures for "IFNγ-dependent metabolic reprogramming restrains an immature, pro-metastatic lymphatic state in melanoma"

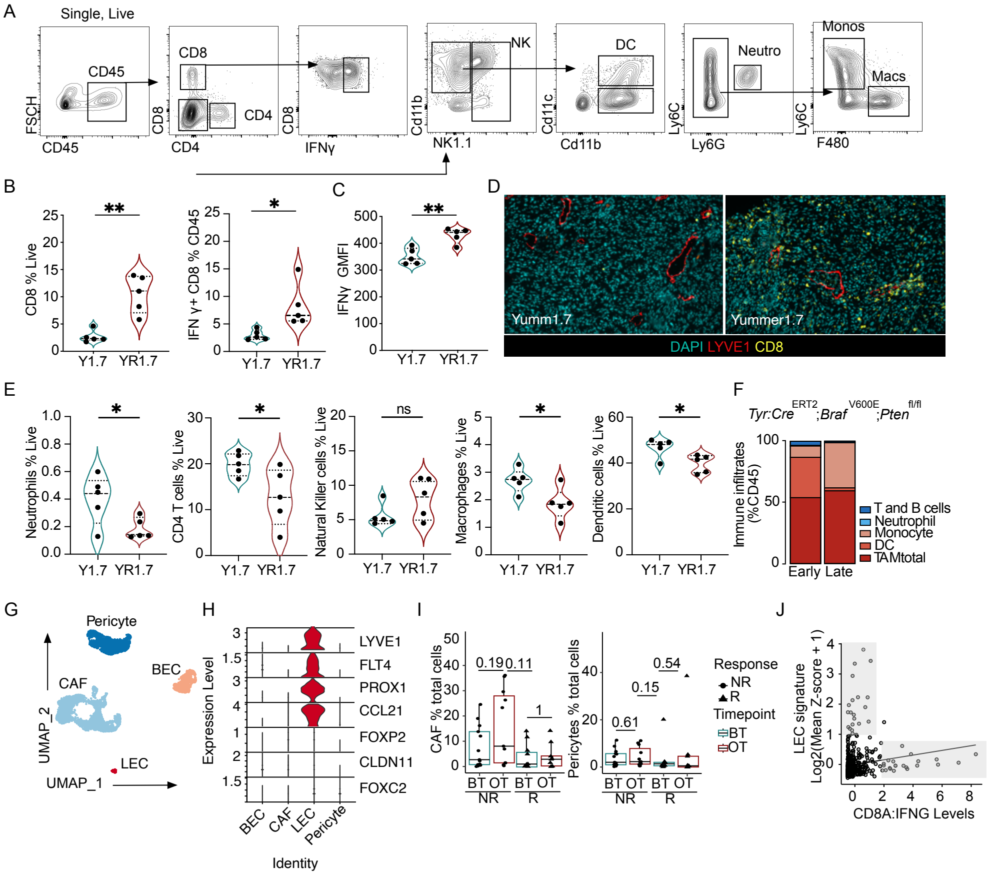


**Supplemental Figure 1. Lymphatic vessels are negatively associated with cytotoxic immune microenvironments.** A. Flow cytometry gating strategy for major immune subsets in YUMM1.7 and YUMMER1.7 tumors. B. $CD8^+$ T cells as a percentage of total live cells each point represents a mouse, (n=5) C. $IFN\gamma^+CD8^+$ T cells as the frequency of $CD45^+$ T cells and C. IFNγ geometrical mean fluorescence intensity (GMFI) on $CD8^+$ T cells, each point represents a mouse (n=5). D. Representative immunofluorescence images of $CD8^+$ T cells (yellow) and LYVE-$1^+$ (red) lymphatic vessels from YUMM1.7 (Y1.7) and YUMMER1.7 (YR1.7) tumors (intradermal, $5x10^5$ cells). DAPI (cyan). Scale bar = 100μm. E. Flow cytometry quantification of immune infiltrates of $CD45^+$, NK cells ($Cd11b^+$, $NK1.1^+$), Macrophages ($CD11b^+$, $F480^+$), Dendritic cells ($CD11c^+$, $MHCII^+$), Neutrophils ($CD11b^+$, $Ly6G^+$), $CD4^+$ and T regulatory ($CD4^+$, $CD25^+$, $Foxp3^+$) cells of the total Live cells in YUMM1.7 and YUMMER1.7 tumors. Each point represents a mouse (n=5). F. Quantification of the immune infiltrates of Tyr::Cre;*Braf* $^{CA/+}$;*Pten* $^{fl/lf}$ tumors at end point as percentage of CD45 cells, G. UMAP plot of the stromal components of the human single cell dataset including cancer-associated fibroblasts (CAFS), blood endothelial cells (BECs), lymphatic endothelial cells (LECs) and pericytes, H. Stacked violin plot of lymphatic endothelial cells markers (LYVE1, FLT4, PROX1, CCL21, FOXP2, CLDN11, FOXC2) enriched in the LEC cluster I. Quantification of CAF and pericytes as a percentage of total cells in non-responder (NR) and responder (R) melanoma patients before and on immunotherapy from the single-cell dataset. J. Scatter plot of LEC and IFNγ$^+$CD8$^+$ T signatures from patient samples from the TCGA dataset, each point is a patient. Unpaired Student's t-tests (B-E), Wilcoxon rank test (I). *$p<0.05$; **$p<0.01$.

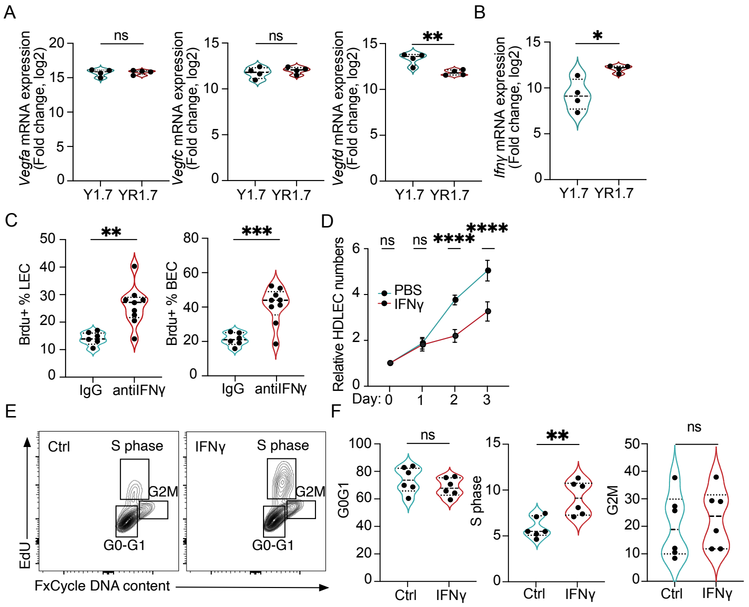


**Supplemental Figure 2. IFNγ puts a cell intrinsic block in LEC proliferation.**

A-B. qPCR analysis of *Ifnγ*, *Vegfa, Vegfc* and *Vegfd* transcripts from Y1.7 and YR1.7 tumor pieces, each point represents a mouse (n=4), C. Quantification of $BrdU^{+}$ LECs and BECs in B16-OVA-VEGFC tumors from IgG or anti-IFNγ treated mice as the frequency of total LECs and BECs, each point represents a mouse (n=6-9), D. Proliferation curve of HDLECs treated with PBS or IFNγ for 3 days, cells were counted daily. E. Representative density flow plots of cell cycle profiles of HDLECs following PBS or IFNγ treatment for 3 days, x axis shows DNA content and y axis EdU incorporation; boxes mark cells in the G0/G1, S, and G2/M phases of the cell cycle, F. Quantification of the the percentages of HDLECs in each cell cycle phase, each points represents a technical replicate across three independent experiments. *$p<0.05$; **$p<0.01$; ***$p<0.001$. Unpaired, student's t test.

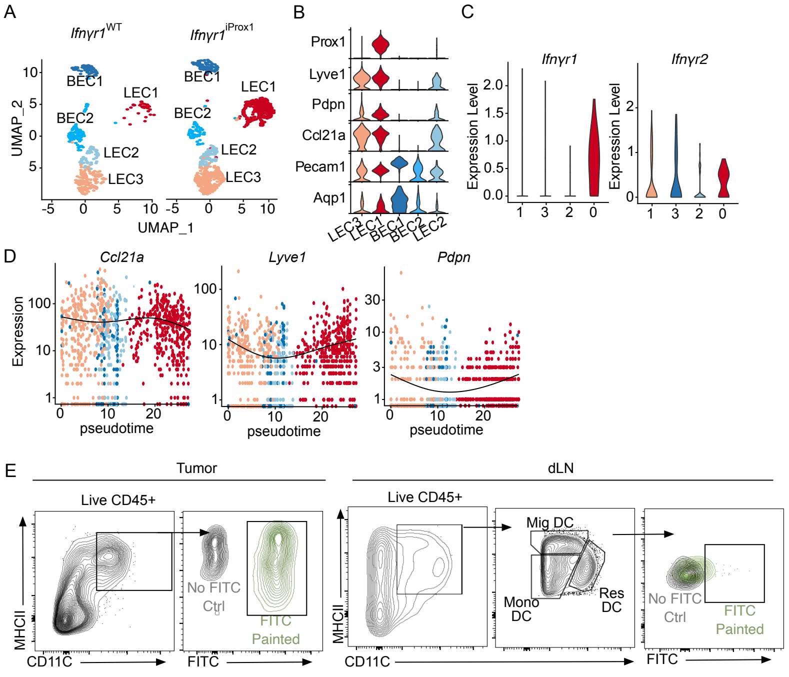


**Supplemental Figure 3. IFNγ constrains an immature LEC state.**
A. Split UMAP plot depicting the LEC and BEC clusters from B16-OVA-VEGFC tumors in *Ifnγr1*$^{WT}$ and *Ifnγr1*$^{iProx1}$ mice, B. Stacked violin plot for LEC and BEC markers, C. Violin plot showing the expression of the *Ifnγr1* and *Ifnγr2* transcripts in the *Ifnγr1*$^{WT}$ mice, D. Gene expression as a function of pseudotime related to figure 3, E. Gating strategy for the quantification of FITC$^{+}$ DCs related to figure 5.

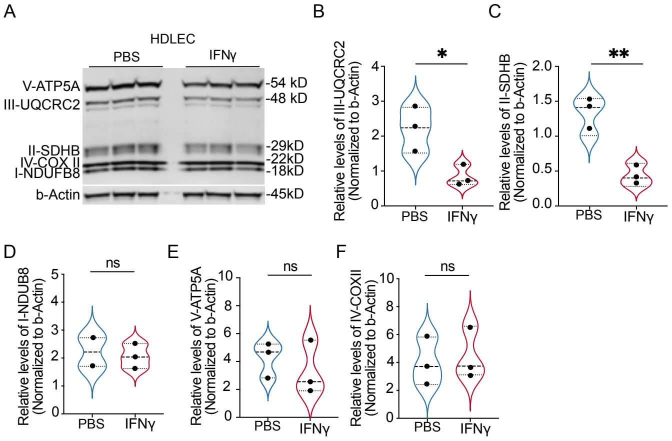


**Supplemental Figure 4. IFNγ downregulates the expression of OXPHOS complexes.**
A. Representative western blot of total OXPHOS proteins and b-actin in HDLEC treated with PBS or IFNγ for 72 hours, B. Quantification of protein normalized to b-actin using ImageJ, every dot is a technical replicate, representative of n=2 biological replicates. *$p<0.05$; **$p<0.01$; ***$p<0.001$. Unpaired, student's t test.

**Table 1.**

| | |
|---|---|
| Tip like (45) | DLL4, ADM, ANKRD37, C1QTNF6, CLDN5, COL4A1, COL4A2, COTL1, EDNRB, FSCN1, GPIHBP1, HSPG2, IGFBP3, INHBB, JUP, KCNE3, KCNJ8, LAMA4, LAMB1, LXN, MARCKSL1, MCAM, MEST, N4BP3, NID2, NOTCH4, PLOD1, PLXND1, PMEPA1, PTN, RAMP3, RBP1, RGCC, RHOC, TRP53ILL |
| Stalk like (45) | ACKR1, AQP1, C1QTNF9, CD36, CSRP2, EHD4, FBLN5, HSPB1, LIGP1, IL6ST, JAM2, LGALS3, LRG1, MEOX2, PLSCR2, SDPR, SELP, SPINT2, TGFBI, TGM2, TMEM176A, TMEM176B, TMEM252, TSPAN7, VWF |
| LEC | LYVE1, PDPN, PROX1, FLT4, PECAM1, CCL21A |
| Petkova et al (44) | PTX3, MRC1, PIK3Ca, CCL2, ACKR2 |
| Notch | DLL4, JAG1, DTX1, JAG2, MAML3, HDAC2, RBPJ, HES1, DTX4, NOTCH3, LFNG, NOTCH4, FZD1, ST3GAL6, DTX1, WNT5A, ARRB1 |
